# A High-Resolution Atlas of the Brain Predicts Lineage and Birth Order Underly Neuronal Identity

**DOI:** 10.1101/2025.06.04.657818

**Authors:** Aaron M. Allen, Megan C. Neville, Tetsuya Nojima, Faredin Alejevski, Devika Agarwal, David Sims, Stephen F. Goodwin

**Author notes:** Co-first author (Aaron M. Allen, Megan C. Neville).

## Abstract

Gene expression shapes the nervous system at every biological level, from molecular and cellular processes defining neuronal identity and function to systems-level wiring and circuit dynamics underlying behaviour. Here, we generate the first high-resolution, single-cell transcriptomic atlas of the adult *Drosophila melanogaster* central brain by integrating multiple datasets, achieving an unprecedented tenfold coverage of every neuron in this complex tissue. We show that a neuron’s genetic identity overwhelmingly reflects its developmental origin, preserving a genetic address based on both lineage and birth order. We reveal foundational rules linking neurogenesis to transcriptional identity and provide a framework for systematically defining neuronal types. This atlas provides a powerful resource for mapping the cellular substrates of behaviour by integrating annotations of hemilineage, cell types/subtypes and molecular signatures of underlying physiological properties. It lays the groundwork for a long-sought bridge between developmental processes and the functional circuits that give rise to behaviour.

## INTRODUCTION

Unravelling the cellular diversity of the nervous system is crucial for understanding how complex behaviours emerge. Although all cells in the nervous system share the same genetic blueprint, differentiation and specialization are driven by differential transcriptional regulation of the genome^1,2^. In *Drosophila melanogaster*’s central brain, subtle transcriptional variations sculpt intricate, highly specialized neural circuits that support complex cognitive functions and behaviours. Mapping the transcriptomes of all neurons within this tissue is essential to uncovering the specific genes and regulatory networks active in each cell type, providing insights into the molecular mechanisms that underlie information processing and adult behaviours.

The adult *Drosophila* central brain develops from approximately 100 pairs of bilaterally symmetric neuronal stem cells, known as neuroblasts, each producing a highly stereotyped lineage of neurons^3–7^. Neuroblasts express unique combinations of transcription factors, creating a "genetic address" that helps define cell types^8,9^. During neurogenesis, neuroblasts generate sibling lineages, or hemilineages, that define anatomical subunits within the brain. The central brain contains two main types of neuroblasts, type I and type II, distinguished by their division patterns and the complexity of their resulting neuronal lineages. Type I neuroblasts generate lineages of approximately 200 neurons, while a small subset of type II neuroblasts (eight per hemisphere) produce larger lineages, averaging over 600 neurons^10,11^. Type II lineages, using intermediate neural progenitor cells, are considered analogous to those in the primate cortex, allowing a single neuroblast to produce many more cells^12,13^.

In both types of neuroblast lineages, tightly regulated temporal and spatial gene expression patterns establish neuronal identities^14,15^. Temporal transcription factors are pivotal in this process, marking distinct stages of neuroblast division and shaping the fate of progeny neurons. While these developmental dynamics are well characterised during embryogenesis and larval stages, their persistence and influence on adult neuronal identity remain less understood. Investigating whether these developmental signatures are retained or modified in the adult brain is essential for understanding the continuity between neurodevelopment and mature brain function^16^.

In this study, we present the most comprehensive single-cell transcriptional atlas of *D. melanogaster* adult central brain neurons to date, generated using single-cell RNA sequencing (scRNA-seq). To sufficiently represent the depth and complexities of cell types within the central brain, we performed a meta-analysis, combining our newly described scRNA-seq data with all publicly available data. This analysis generated an atlas of 329,466 neurons, representing an average 9.8x depth of coverage of every single neuron of the central brain.

Our findings reveal that adult central brain neurons are largely transcriptionally defined by their developmental histories. Neuroblast hemilineages emerge as the primary genetic units that define transcriptionally distinct neuronal cell types in the adult central brain. Within these hemilineages, we observe that transcriptional signatures linked to different birth order windows are maintained into adulthood. Notably, early-born neurons exhibit distinct transcriptional profiles compared to late-born neurons, uncovering a novel axis of neuronal diversity in the adult brain that likely reflects differing developmental demands.

Variation in gene expression drives plasticity, learning, and adaptation, while more drastic gene disruptions contribute to neurological disorders; thus, understanding the underlying genetic mechanisms active in distinct neurons within the brain is essential to linking molecular processes to circuit function. Moreover, leveraging the distinct molecular signatures of neuronal subtypes enables targeted genetic access, facilitating the causal investigation of circuit dynamics. Our high- resolution single-cell transcriptomic atlas of the adult *Drosophila* central brain represents a significant advancement in neurogenomics. By providing an unprecedented level of resolution and integrating genetic, developmental, and anatomical data, this resource provides a comprehensive framework for dissecting the molecular logic of neuronal diversity. More broadly, our findings offer a platform for comparative studies across species, informing general principles of brain development, organization and evolution.

## RESULTS

### Generating a high-depth single-cell transcriptomic atlas of the adult central brain

To capture the transcriptional diversity of neuronal cell types within the adult *Drosophila* central brain, we first generated a sexed single-cell RNA-seq dataset, excluding the visual system, which yielded ∼1.6× coverage of central brain neurons (Figure 1A; Figure S1). Given the complexity of neuronal cell types in this tissue, we concluded that significantly greater coverage was required to refine neural identity. We therefore integrated multiple publicly available datasets, all generated using 10x Chromium 3’ chemistries (v2 and v3) from single-cell (sc) and single-nucleus (sn) preparations. These datasets spanned diverse tissues, including central brain-enriched dissections, whole brain, optic lobes, retina, and whole heads, and together formed a meta-atlas of >1 million cells/nuclei from the adult *Drosophila* head (Figure S2A-B).

**Figure 1.**
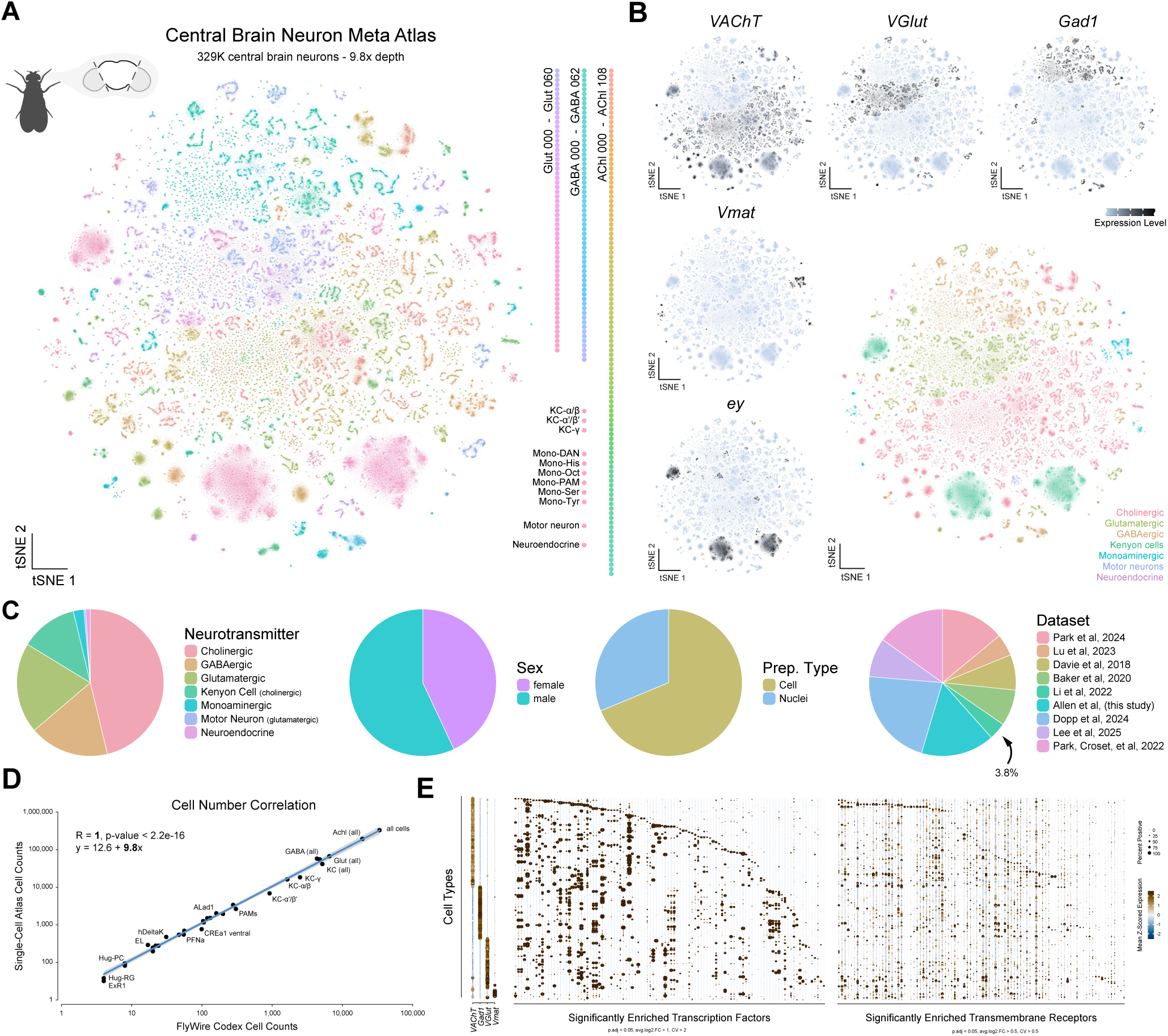
As single-cell meta-atlas of *D. melanogaster* central brain neurons. (A) t-distributed stochastic neighbour embedding (t-SNE) of 329,466 central brain neurons representing a 9.8x cellular depth of coverage, coloured by cell type. (B) Broadly defined neural cell type across the central brain. t-SNEs showing the distribution of cells expressing biomarkers for the fast-acting neurotransmitters acetylcholine (*VAChT*), glutamate (*VGlut*) and GABA (*Gad1*), as well as monoamines (*Vmat*), and Kenyon cells (*ey*). Expression shown in black, intensity is proportional to the log-normalised expression levels. Coloured t-SNE based on assigned cluster identities. (C) Pie charts representing the cellular composition of the central brain neuron atlas based on assigned broad cell type, sex, preparation type (single-cell vs single-nuclei), and dataset origin. (D) Correlation between single-cell atlas-derived cell counts and FlyWire-derived cell counts, demonstrating strong agreement (R = 1, p-value < 2.2e-16). Each point represents an anatomically defined neuronal cell type. (E) Dot plot of the expression of neurotransmitter identity (left), significantly enriched transcription factors (middle) and transmembrane receptors (right) across central brain neuronal cell types.

We refined this atlas in three steps (Figure S2C-G). First, we annotated and removed non-neuronal cells, yielding a ∼700,000 neuronal head meta-atlas (Figure S2D). Next, we identified and excluded optic lobe and peripheral neurons (Figure S2E-F). The remaining central brain neurons were re-clustered, resulting in a high-resolution transcriptomic atlas of 329,466 central brain neurons, achieving an unprecedented 9.8x average depth of coverage of every neuron in the central brain (Figure 1A; Figure S2G; Figure S3A-E).

Using established marker genes, we assigned broad cellular identities to 246 transcriptionally distinct neuronal clusters (Figure 1B). The proportional makeup of broad cell types was consistent with previous reports^17–19^ (Figure 1C; Figure S3A). This atlas has robust coverage of both sexes (Figure 1C), and sex differences in the central brain are examined in detail in a companion study^20^. Notably, central brain neurons from the Fly Cell Atlas (FCA) comprise only 3.8% of the neurons recovered in our dataset^21^. In the FCA dataset, most central brain neurons remained unresolvable and therefore unannotated^22^, indicating that the central brain’s complexity required far greater coverage. Indeed, no individual dataset used here was sufficient on its own to describe this complexity. Thus, our cost-effective approach is a critical addition to existing atlases.

To estimate the cellular depth of coverage of our atlas, we compared the cell counts of our annotations with anatomical estimates from the Flywire connectome^10,11^. These annotations ranged from scarce populations of four neurons per brain (e.g., Hug-RG or ExR1) to all ∼34K neurons of the central brain and achieved a depth of coverage of 9.8x with a correlation R = 1 (Figure 1D). Gene ontology analysis of cell type-defining genes (Figure S3F) found transcription factors and transmembrane receptors define much of the transcriptional heterogeneity of neuronal cell types within the central brain (Figure 1E). Indeed, unique combinations of transcription factors can be leveraged to define the complexity of cell types across the central brain (Figure 1E; Figure 2).

**Figure 2.**
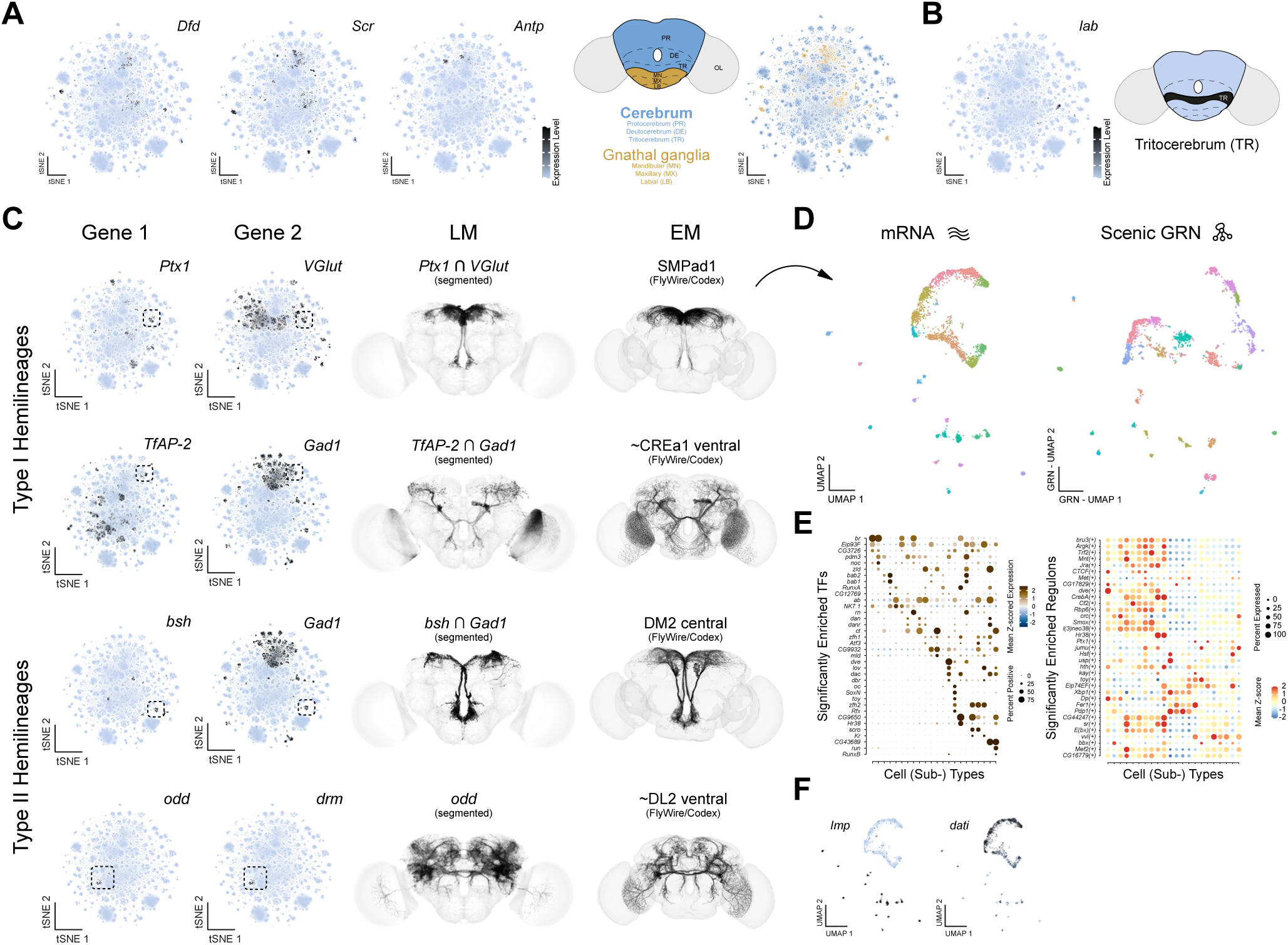
Hemilineages define transcriptional cell types in the adult central brain. (A) t-SNEs of Hox gene expression (*Dfd*, *Scr*, and *Antp*) associated with the gnathal ganglia across the central brain. Schematic (right) representing regions along the n-anterior/n-posterior neuraxis associated with the cerebrum and the gnathal ganglia. Cells assigned in t-SNE based on differential Hox gene expression: cerebrum in blue and gnathal ganglia in gold. (B) t-SNEs of *lab* gene expression (left) associated with the tritocerebrum with schematic (right). (C) Genetic intersections identify anatomically distinct cell types as hemilineages. t-SNEs showing gene pairs (*Ptx1* and *VGlut*, *TfAP-2* and *Gad1*, *bsh* and *Gad1*, *odd* and *drm*) used to identify cell types (left). Light microscopy (LM) images (middle) showing neuronal populations identified via intersecting gene pairs or individual gene reporter (*odd*). Electron microscopy (EM) reconstructions of selected hemilineages from FlyWire dataset (SMPad1, CREa1 ventral, DM2 central, DL2 ventral – tildes denote estimations, see Figure S4) based on transcriptionally defined cell types. LM images have been segmented; full expression patterns are shown in Figure S4. For full genotypes, see STAR Methods. (D) Uniform manifold approximation and projections (UMAPs) of the SMPad1 hemilineage subclustered based on gene expression (left) and SCENIC GRN analysis (right), revealing shared and distinct regulatory programs across SMPad1 subtypes. (E) Dot plot of the expression of significantly enriched transcription factors (left) and regulons (right) across SMPad1 hemilineage subtypes (below). (F) UMAPs of *Imp* (enriched in early-born neurons), and *dati* (enriched in late-born neurons) across SMPad1 hemilineage subtypes.

### Transcriptionally defined cell types in the adult brain reflect developmental origins

The *Drosophila* central brain encompasses the cerebrum and the gnathal ganglia. Each consists of three neuromeres, the morphological units along the body axis^23^. Previous work, including ours, has shown that the Hox homeodomain transcription factors, which define neuromeres, maintain their regional expression in the adult^24,25^. We used expression from members of the Antennapedia-complex of Hox genes to define cell types associated with the gnathal ganglia; while cerebrum-associated cell types were identified based on the absence of Hox gene expression, consistent with the anterior CNS being largely Hox-negative^26,27^ (Figure 2A-B). In the VNC, serially homologous cell types represent a conserved pattern of neuronal organization across segments^28^. The extent to which cell types are repeated or homologous within the brain and between the brain and VNC is unclear. Anatomically comparing neuronal types and circuits to identify homologous structures is challenging due to the high degree of specialisation in these two tissues. Our analysis suggests that leveraging single-cell transcriptomics will substantially advance future efforts to identify serially homologous cell types across the central nervous system.

To determine the anatomical identity of transcriptionally defined cell types, we applied an intersectional genetic strategy (see STAR Methods). We anatomically characterised the genetic intersection of two genes co-expressed in a given transcriptionally defined cell type, then matched with cell types based on morphology within the adult connectome^10,11^. Using this approach, we repeatedly and consistently (8/8) identified intersected cell types representing entire hemilineages^20,29^ (Figure 2C; Figure S4), just as we and others had previously seen in the VNC^24,30^. For example, the co-expression of the transcription factor *Ptx1* and the glutamatergic neuron marker *VGlut* corresponded to hemilineage SMPad1 (Figure 2C, top; Figure S4B-F). Thus, the overarching transcriptional relationship between neurons within the adult central brain appears to reflect their shared developmental origins.

As cell types within hemilineages are anatomically diverse, we assessed their transcriptional heterogeneity. By way of example, subclustering analysis of the SMPad1 hemilineage revealed 22 transcriptionally distinct subtypes (Figure 2D). Subtype-specific transcription factors were identified, and gene regulatory networks (GRNs) were inferred using SCENIC^31,32^ (Figure 2E). Sub-clustering based on SCENIC-defined GRNs closely matched clustering based on gene expression alone, with regulons (transcription factors and their putative targets) defining hemilineage subtypes (Figure 2D-E; Figure S5). Similarly, by sub-clustering all transcriptionally defined neuronal cell types in the central brain (Figure 1A), we can resolve 4,167 distinct neuronal subtypes offering an unprecedented view of adult brain cellular diversity (Figure S6). Ultimately, transcriptional identity and GRNs can be integrated with neuronal connectivity maps to reveal how molecular programs shape circuit architecture and drive behaviour.

The most striking observation from our hemilineage analyses was the distinct spatial arrangements of cellular subtypes in our UMAP, reflecting the transcriptional relationships amongst subtypes. Some subtypes formed discrete, compact groupings resembling "punctate" clusters, representing cells with highly similar gene expression profiles. In contrast, other subtypes appeared to group together, forming elongated "serpentine" clusters, reflecting a transcriptional continuum between related subtypes. We examined their associated gene expression profiles to understand the molecular basis of these distinct cluster morphologies and identified the neurodevelopmental gene *Imp* as highly enriched in subtypes exhibiting punctate morphology, while *dati* expression was highly correlated with serpentine subtypes (Figure 2F). *Imp* and *dati* are part of a transcriptional temporal patterning cascade driving neurogenesis in the *Drosophila* CNS, with *Imp* associated with early-born neurons and *dati* associated with late-born neurons^24,33–36^. Thus, the transcriptional relationships between punctate and serpentine subtypes appear to reflect neuronal birth order within hemilineages of the adult central brain.

### Neurodevelopmental gene expression distinguishes neuronal birth order across central brain subtypes

To explore the broader relevance of birth order, we analysed the relationship between *Imp* and *dati* expression across all neurons in the central brain. As seen in the VNC^24^, their expression was largely mutually exclusive (Figure 3A-B). Notably, optic lobe-derived neurons lacked expression of both genes, suggesting region-specific developmental programs (data not shown). *Imp*-enriched neurons accounted for approximately 30% of central brain neurons and consistently exhibited higher levels of total transcripts and total genes (Figure S8C-E). These transcriptional differences suggest inherent distinctions in neuronal populations in the adult based on birth order.

**Figure 3.**
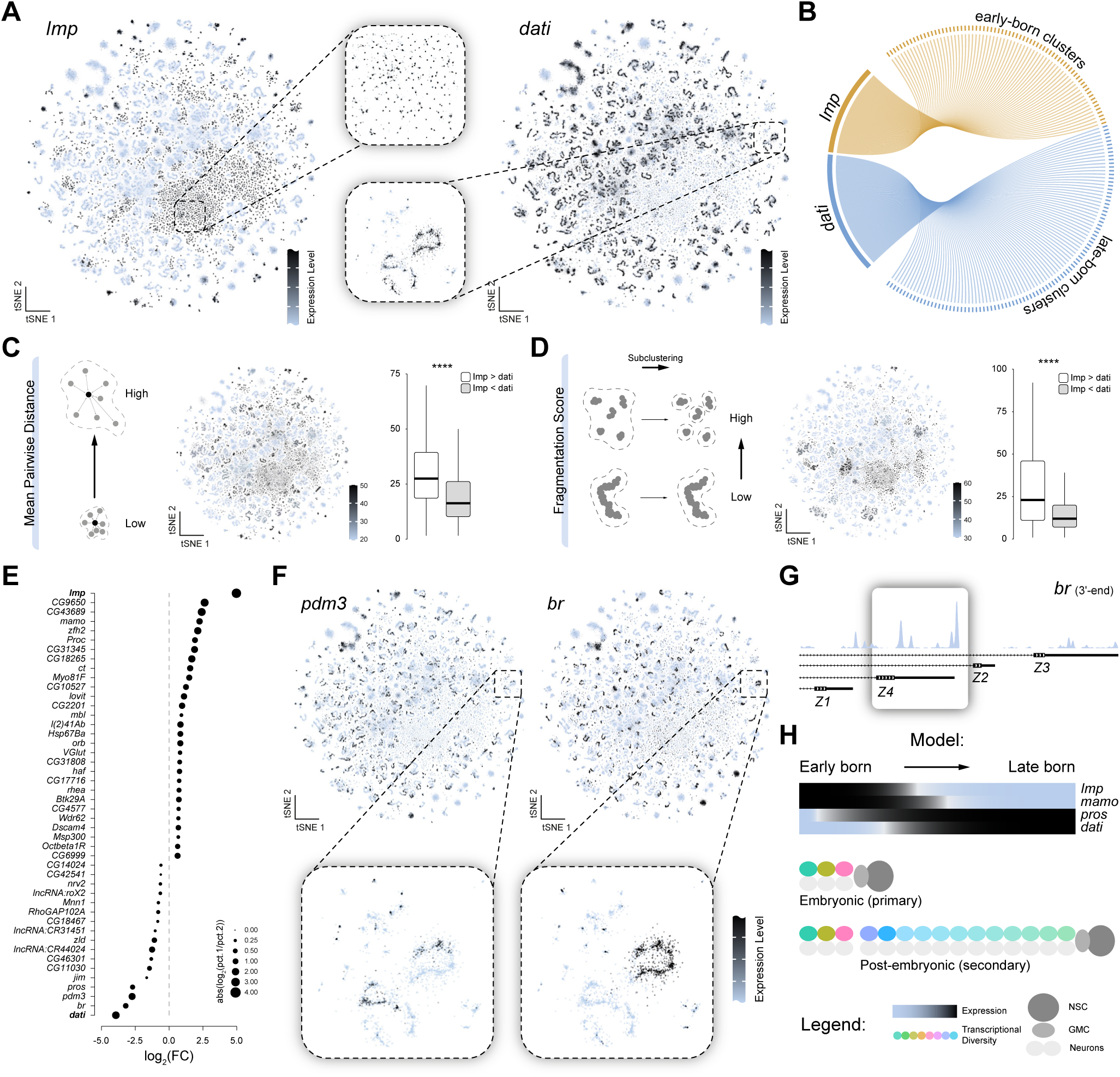
Temporal patterning and transcriptional diversity in the central brain. (A) t-SNEs showing the expression of *Imp* (left) and *dati* (right) across all central brain neurons (Kenyon cells removed). Insets (middle) highlight distinct clustering of *Imp*- and *dati*-expressing cells, reflecting their transcriptional and spatial organization. (B) Chord diagram of *Imp* and *dati*, and the clusters for which they are significantly enriched, illustrating their mutual exclusivity. (C) Schematic (left) of average pairwise distances among cells within a cluster (dashed line), where high distances reflect greater transcriptional diversity. t-SNE (middle) coloured by average pairwise distances by cluster. Box plot of average pairwise distance across clusters split by *Imp+* neurons compared to *dati+* neurons (Bonferroni corrected Wilcoxon Signed-Rank Test, ****p < 0.0001). (D) Fragmentation score analysis comparing cluster cohesiveness between *Imp*+ and *dati+* neurons. Schematic (left) showing that high fragmentation scores reflect more fragmented clusters (clusters within dashed lines). t-SNE (middle) coloured by fragmentation scores. Box plot (right) showing significantly higher fragmentation scores in *Imp*+ neurons (Bonferroni corrected Wilcoxon Signed-Rank Test, ****p < 0.0001). (E) Ranked dot plot of genes differentially expressed between *Imp*+ and *dati*+ neurons. Dot size represents the relative specificity of each gene’s expression in *Imp*+ vs. *dati*+ neurons. (F) t-SNEs (top) showing the expression of neurodevelopmental transcription factors *pdm3* (left) and *br* (right) across all central brain neurons showing enrichment in distinct neuronal subclusters. Insets (below) highlight the specific localisation of these transcription factors within transcriptionally distinct regions of presumed hemilineages. (G) Isoform-specific expression (top) of *br* (Z1–Z4) within adult central brain neurons, showing high levels of Z4 isoform expression. (H) Schematic model representing how temporal patterning during *Drosophila* neurogenesis (top) shapes transcriptional organisation of neurons in the central brain. *Imp* marks early-born neurons with higher transcriptional diversity, while *dati* is enriched in late-born neurons. NSC (neuronal stem cell), GMC (ganglion mother cell).

To determine whether these differences reflect biological heterogeneity or technical artifacts, we employed three separate analytical strategies: (1) varying the number of principal components used in dimensionality reduction, (2) projecting early- vs late-born into a shared embedding, and (3) independently subclustering each population. When we varied the number of principal components, *Imp* and *dati* remained mutually exclusive, but the distinct cluster morphologies only emerged as we increased the number of principal components, and indeed *Imp* and *dati* contribute to many principal components (Figure S7A). Projecting early- vs late-born into a shared embedding maintained both the mutual exclusivity and the punctate versus serpentine t-SNE morphologies (Figure S7B). Finally, independently subclustering early-born and late-born neurons also preserved their respective t-SNE morphologies (Figure S7C). These findings suggest that the differences in t-SNE morphologies are intrinsic and robust features of the developmental stage-specific transcriptional complexity of early-born and late-born neurons rather than artifacts of dimensionality reduction.

To quantify transcriptional relationships within clusters, we calculated average pairwise distances, fragmentation scores, and modularity scores for cells expressing *Imp* versus *dati* (Figure 3C-D; Figure S8F-I). Early-born, *Imp*-enriched cells were more dispersed and fragmented, leading to their characteristic punctate t-SNE-morphology. In contrast, late-born, *dati*-enriched cells exhibited more connected transcriptional relationships, consistent with their serpentine t-SNE- morphology. Our findings reveal that *Imp* and *dati* not only mark neuronal birth order but also correlate with fundamentally distinct transcriptional profiles in the central brain.

To further explore differences between these neuronal populations, we identified genes most enriched in early- versus late-born neurons. This analysis uncovered several neurodevelopmental genes indicative of neuronal birth order strongly associated with *dati* expression (Figure 3E). Notably, the transcription factors *Pdm3* and *br* were the most enriched in *dati*+ cells, with both genes displaying regional expression across multiple serpentine clusters in our atlas (Figure 3F; Figure S8J). A specific isoform of *br*, Z4, known to mark cells born during larval L2-L3 ecdysis^37^, was the predominant isoform expressed in our atlas (Figure 3G). Cells expressing *br-Z4* consistently mark cells born at a specific developmental window across hemilineages, a pattern evident in our data. These observations support our broader model, which proposes that transcriptionally defined cell types represent hemilineages while subtypes reflect birth order. Moreover, within a hemilineage, early-born subtypes are more transcriptionally distinct from each other and from late-born subtypes, suggesting that neurogenic timing not only shapes identity but also influences the granularity of transcriptional diversity within lineages (Figure 3H).

### Systematic reconstruction of birth order reveals repeated temporal transcription factor programs

Our findings suggest that transcriptional profiles in the adult brain retain signatures of developmental timing, enabling systematic reconstruction of neuronal birth order across hemilineages. As proof of principle, we investigated the anterodorsal antennal lobe (AL) olfactory projection neuron hemilineage, ALad1, whose birth order and transcriptional identities have been largely established^38,39^. We identified ALad1 neurons in our atlas, re-clustered, and transferred previously defined subtype annotations (Figure 4A-C; Figure S9; see STAR Methods). To investigate transcriptional temporal dynamics within the hemilineage, we performed pseudotime analysis, which orders cells along a continuous trajectory based on their gene expression profiles, anchoring the earliest-born neurons to *Imp* expression. This trajectory strongly correlated with experimentally validated birth order (Figure 4D), confirming that pseudotime recapitulates temporal lineage progression in mature neurons and can be used to identify birth order-associated transcriptional programs (Figure 4E).

**Figure 4.**
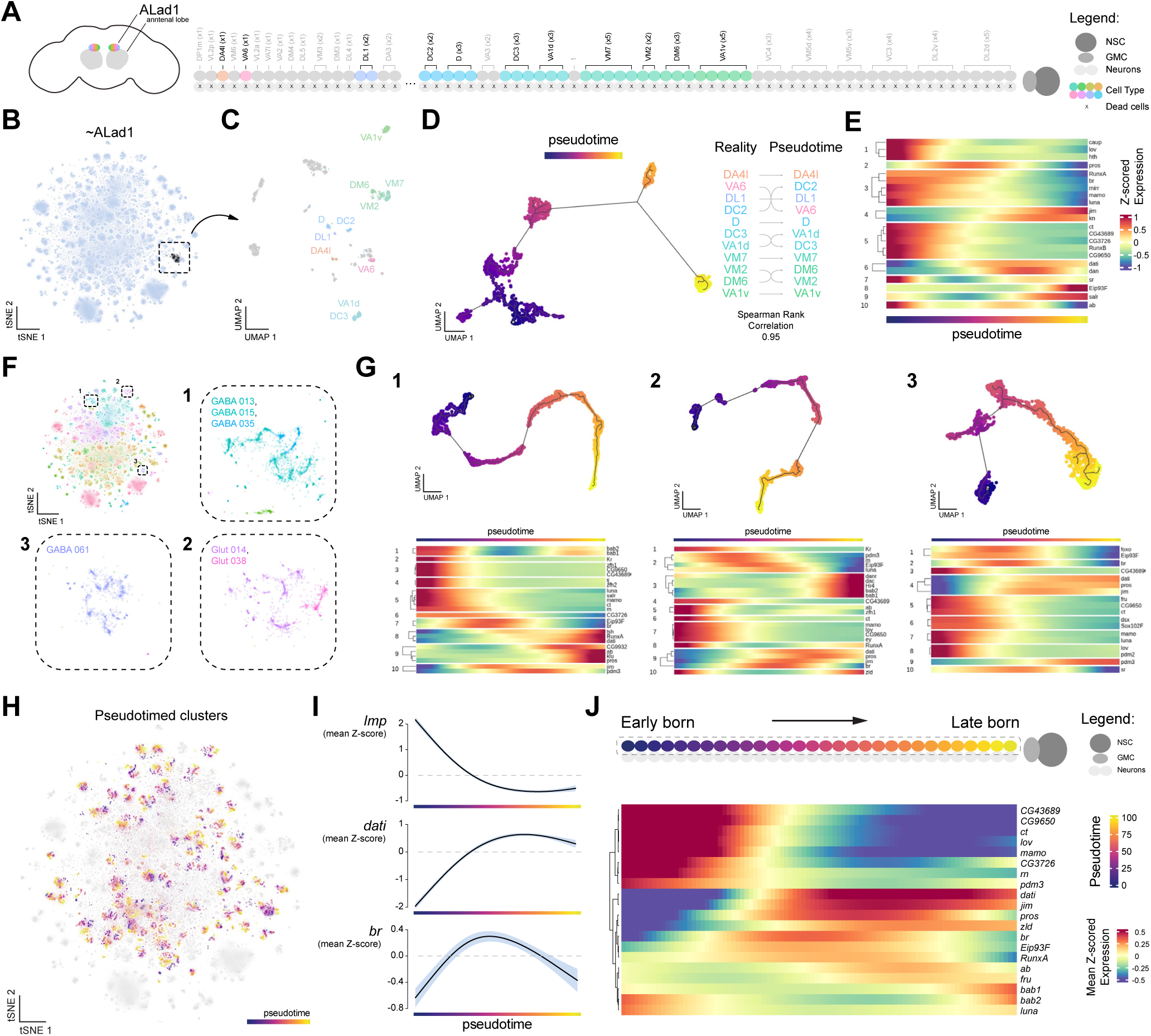
A common set of repeated transcription factors defines transcriptional subtypes within hemilineages. (A) Schematic of the ALad1 hemilineage and its associated subtypes, arranged in known developmental order (top), dead neurons of the sister hemilineage are shown below. (B) t-SNE highlighting ALad1 neuronal cluster (in black) extracted from the central brain atlas. (C) UMAP of ALad1 hemilineage, with previously described transcriptionally defined subtypes annotated (see STAR Methods). (D) Pseudotime trajectory (left) compared to the real birth order of ALad1 cell types (right). A strong correlation (Spearman rank correlation = 0.95) demonstrates that pseudotime accurately recapitulates temporal cell type progression. (E) Heatmap showing gene expression dynamics of transcriptional markers along pseudotime, with key transcription factors defining early, middle, and late birth order identities within the hemilineage. (F) t-SNE of central brain atlas highlighting three distinct presumed hemilineage cell types for pseudotime analysis (1, 2, and 3) with zoomed-in regions. (G) Pseudotime trajectories (top) for the three highlighted regions representing presumed hemilineage. Heatmaps (bottom) showing gene expression changes of significantly enriched transcription factor across pseudotime. (H) t-SNE of pseudotime-annotated neuronal clusters across the entire central brain, coloured by pseudotime progression per cluster. (I) Line plots showing mean z-scored expression of key temporal markers (*Imp*, *br*, *dati*) along the pseudotime axis, capturing their expected early, middle, and late transcriptional dynamics, respectively. (J) Pseudotime across all clusters identify novel genetic markers of neurodevelopmental windows that repeat across cell types. Heatmap averaging pseudotime-ordered gene expression across central brain clusters, with known and novel transcription factors found to broadly represent subtypes of cells within hemilineages.

Given that unsupervised clustering in our atlas largely reflects hemilineage identity (Figures 2-3), we expanded pseudotime analysis to include multiple hemilineages (Figure 4F-G). We identified several temporal transcription factors that are repeatedly utilised across these selected hemilineages (such as *bab1/bab2*, *dan/danr*, *Eip93F*, and *zld*), reflecting conserved temporal developmental programs. To systematically examine birth order-associated transcriptional dynamics, we extracted, re-clustered, and applied pseudotime analysis across the majority of *dati*+ hemilineages (Figure 4H; see STAR Methods). By examining the genes that vary across pseudotime within presumed hemilineages, we can explore repetitive transcriptional signatures across the brain. Averaging the expression of the genes that varied over the pseudotime confirmed the expected temporal expression patterns of *Imp*, *br*, and *dati*, representing early-, mid-, and late-born neurons, respectively (Figure 4I). Filtering for transcription factors with repeat occurrences across presumed hemilineages revealed known and novel temporal regulators used throughout the central brain, highlighting shared mechanisms underlying neuronal birth order^9,14,37,40,41^ (Figure 4J). These findings indicate that temporal patterning mechanisms governing neurogenesis are deeply embedded in the adult transcriptional landscape across the central brain. Furthermore, these repeated temporal regulators represent an orthogonal axis to the gene programs underlying hemilineage in defining neuronal identity.

### Uncovering physiological properties through cell type annotation of the central brain

To further resolve the molecular and physiological identity of neurons in the adult central brain, we annotated neuronal cell types utilising existing single-cell and bulk RNA-seq data sets. Neurons in the central brain are primarily derived from one of three types of neuroblasts – type I, type II, and mushroom body (MB) (Figure 5A; reviewed in^12,42–44^). To distinguish lineage origin, we reprocessed and integrated published larval type II scRNA-seq data^45,46^ with our adult atlas (Figure 5B), enabling the systematic assignment of adult neurons to type I or type II neuroblast lineages (Figure 5C). This analysis reinforced the lineage-based organisation of transcriptionally defined cell types and confirmed that the previously identified hemilineages correspond to their expected neuroblast types (Figure 2B). We re-clustered all type II neurons, generating an adult type II neuroblast lineage atlas (Figure 5D).

**Figure 5.**
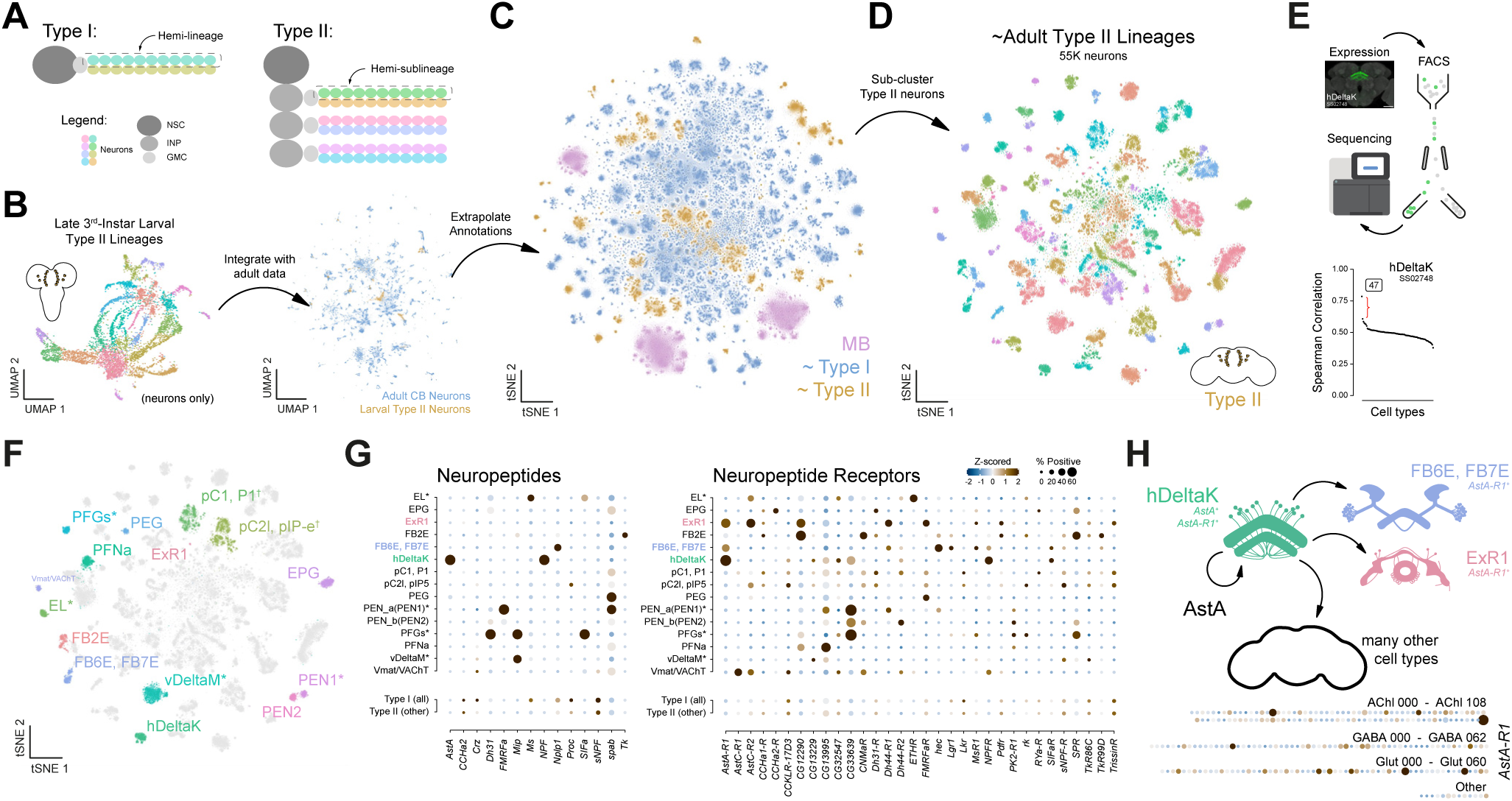
Integrative analysis of type II lineages in the central brain. (I) Schematic illustrating type I hemilineages and type II hemi-sublineages. (J) UMAPs of larval type II data (left) integrated with adult central brain data (right). Reprocessed larval type II data originally from Michki et al., 2021^45^ and Rajan et al., 2023^46^. (K) t-SNE of the central brain neuronal atlas with extrapolated annotations highlighting three neuroblast lineage types – type I (blue), type II (gold), and mushroom body (MB, pink). (L) t-SNE of ∼55,000 type II neurons subclustered, representing 98 cell types. (M) Workflow for genetic labelling (GFP+ cells) and transcriptional analysis of hDeltaK neurons: targeted expression based on genetic access, FAC sorting, and RNA sequencing (top). Spearman correlation analysis of cell types in the adult type II atlas compared to hDeltaK sequencing data (bottom). Cluster 47 shows the highest correlation. Immunostaining of hDeltaK (SS02748) obtained from FlyLight. Bulk RNA-seq data originally Wolf et al., 2025^49^. (N) Annotated t-SNE of type II atlas (see Figure S10-S11, STAR Methods). (O) Dot plot of neuropeptide (left) and neuropeptide receptor (right) expression across annotated type II cell types, revealing potential signalling interactions. (P) Schematic of AstA signalling in hDeltaK neurons and AstA-R1 expressing cell types. hDeltaK neurons (*AstA*+, *AstA-R1*+) and other *AstA-R1*+ central brain cell types (e.g., FB6E, FB7E, ExR1) likely represent localised AstA-modulated circuits. A dot plot of *AstA-R1* expression across the central brain reveals a broader network of potential signalling partners.

To map transcriptionally defined clusters in our type II atlas to anatomically defined cell types, we leveraged previously generated bulk RNA-seq datasets of genetically labelled and fluorescence-activated cell-sorted central brain populations^47–49^ (Figure 5E; Figure S10). Correlating these profiles with our atlas (see STAR Methods) enabled high-confidence annotation of type II cell types based on distinct transcriptional signatures (Figure 5F; Figure S10). Additionally, we leveraged genetic intersection^20^ and neuropeptide (NP) expression as molecular markers^49^ to identify central brain cell types, resulting in the most comprehensive molecular map of type II-derived cell types in the adult brain to date (Figure 5G; Figure S11).

To highlight the functional relevance of our annotations, we examined NP and NP-receptor expression across the type II atlas (Figure 5G-H; Figure S11D-E). In agreement with prior studies^49^, *AstA* expression was enriched in hDeltaK neurons, while its receptor, *AstA-R1*, was detected in FB6E, FB7E, ExR1, and hDeltaK neurons, suggesting local and autoregulatory *AstA* signalling within the central complex, a hub for navigation and behavioral state control^50,51^ (Figure 5G–H). Interestingly, hDeltaK neurons had the highest expression of *AstA* and *AstA-R1* amongst all cell types across the central brain, suggesting that autocrine signalling within this population facilitates a homeostatic feedback loop (Figure 5H; Figure S12A). Additionally, we identified multiple unannotated *AstA-R1*+ cell types distributed across the central brain, highlighting new potential targets of *AstA* signalling and expanded roles for hDeltaK neurons in modulating both local and long-range circuits (Figure 5H, bottom). Our atlas provides a framework for targeted genetic access to these neuronal populations and enables mechanistic dissection of neuropeptidergic regulation of central complex-dependent behaviours such as navigation or arousal-related activity states.

### Transcriptional and functional specialization of neuroendocrine cells in the central brain

By and large, cells cluster by hemilineage when performing unsupervised clustering (grouping cells based on their gene expression patterns), reflecting their shared developmental origins. However, this was not always the case, as some cells cluster due to convergent gene expression patterns reflecting shared physiological properties. This pattern is especially evident in neuroendocrine neurons, which secrete neuropeptides to regulate homeostatic and behavioural processes in response to internal state and environmental cues^52^. As shown previously^24^, the transcription factor *dimm*, essential for the differentiation of neurosecretory cells^53–55^, enables robust identification of neuroendocrine populations in the adult brain (Figure 6A). Isolating *dimm*+ cells and sub-clustering based on gene expression and GRNs revealed 34 highly transcriptionally distinct neuroendocrine subtypes (Figure 6B-C). Consistent with their embryonic origin, all neuroendocrine subtypes expressed *Imp*, further supporting its role as a marker for early-born neurons (Figure 3A). Similarly, despite their distinct developmental origins, monoaminergic cells clustered together within our atlas, as detailed in Figure S13.

**Figure 6.**
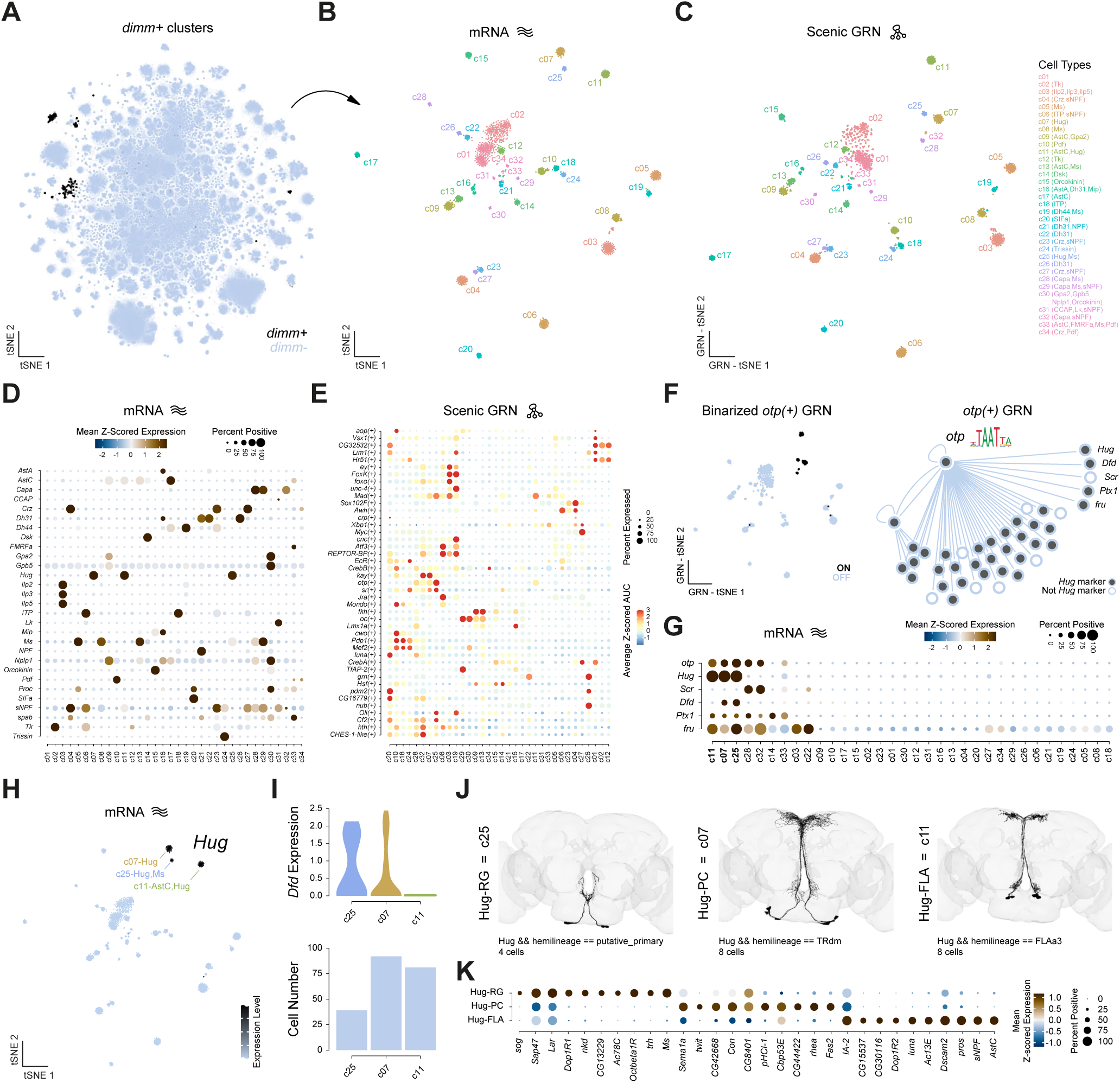
Transcriptional diversity of central brain neuroendocrine cell types. (A) t-SNE highlighting neuropeptide expressing cell types based on the expression of the neurosecretory cells specific transcription factor *dimm*. (B) t-SNE of subclustered neuroendocrine cell types, showing transcriptionally distinct cell types. (C) GRN-t-SNE of subclustered neuroendocrine cells based on SCENIC GRN analysis, identifying regulons driving neuroendocrine specialisation. (D) Dot plot of neuropeptide expression across neuroendocrine subtypes. (E) Dot plot of significantly enriched transcription factor regulons identified by SCENIC GRN analysis across neuroendocrine subtypes. (F) Binarized regulon activity of the transcription factor *otp* in neuroendocrine subtypes (left). GRN showing *otp* driving the expression of multiple genetic markers for *Hug*+ cell types (right). (G) Dot plot of *otp*-regulated genes (*Hug*, *Scr*, *Dfd*, *Ptx1*, and *fru*) across neuroendocrine subtypes, showing enriched expression of *Hug* and *otp* in three subtypes (c07, c11, c25). (H) t-SNE showing *Hug* expression in neuroendocrine subtypes, highlighting *Hug*-expressing subtypes with other co-expressed NPs. (I) Violin plots of *Dfd* expression levels across *Hug*-expressing subclusters (top) and bar plot showing the number of cells in each subcluster (bottom). (J) EM reconstructions of the three *Hug*-expressing neuroendocrine subtypes in the central brain from the FlyWire dataset. Anatomical and transcriptional subtypes have been correlated based on *Dfd* expression and cell number representation (RG = c25, PC = c07, FLA = c11). (K) Dot plot of the top genes defining *Hug*+ subtypes.

To probe the functional diversity of neuroendocrine subtypes, we examined NP expression, associating specific NPs with each subtype (Figure 6D). Over half of the neuroendocrine subtypes showed weak or no expression of fast-acting neurotransmitter or monoaminergic neuron markers, suggesting they are specialised for NP release (Figure S12). Despite their shared functional role in neuropeptide release, neuroendocrine cell types arise from different hemilineages, giving each subtype distinct transcriptional signatures. Moreover, SCENIC-based regulon analysis further confirmed broad regulatory heterogeneity, revealing diverse GRN usage across subtypes (Figure 6E). While neuroendocrine neurons constitute a small fraction of the central brain, many additional neurons express NPs at lower levels, likely supporting local signalling^56^. We classified NPs into four categories based on these expression patterns – very broad, broad, restricted, and very restricted – revealing that some NPs show broad expression within, but restricted to, specific neurotransmitter cell types (e.g., *spap* with acetylcholine and *Nplp1* with glutamate/GABA). In contrast, others (e.g., *Orcokinin* and *Ilp2*) are restricted to a few neuroendocrine neurons (Figure S12). These findings highlight NP-signalling’s transcriptional and functional diversity in the central brain, reflecting their specialised roles in coordinating physiological and neuronal processes.

To demonstrate that our atlas reliably links transcriptional identity to anatomical identity, even for rare cell types, we focused on neuroendocrine subtypes expressing the neuropeptide *Hugin* (*Hug*), which have been shown to act as integrators of internal state and sensory cues, modulating behaviours such as feeding, locomotion, and circadian activity^57–61^. We focused on the homeodomain transcription factor *otp* and its regulon, since *otp* is predicted to bind *Hug*, and its regulon showed restricted expression across neuroendocrine subtypes, particularly in three *Hug*- expressing subtypes (Figure 6F-G). Most genes identified within the *otp* regulon were significantly enriched as markers for all *Hug*-expressing subtypes (Figure 6F). Using a combination of cell number representation and HOX gene expression (e.g., *Dfd*), we linked transcriptional subtypes to anatomically defined subtypes in Flywire (Figure 6H-J). By examining specifically enriched genes for different *Hug*-expressing subtypes (Figure 6K), we uncovered new insights into their functional properties, including the co-release of other NPs and their potential to respond to distinct monoamines (e.g., *Octbeta1R* in Hug-RG neurons). Importantly, these neurons constitute rare cell types – four cells per brain – demonstrating that our atlas possesses sufficient resolution to identify unique transcriptional profiles and functional characteristics of rare neuronal populations. As the necessity for cell-specific genetic access becomes increasingly important, our atlas serves as a valuable resource for disentangling neuronal heterogeneity present in existing tools, even for rare cell types, enabling more precise and interpretable behavioural studies in the future^62^.

### An interactive web resource for exploring *Drosophila* central brain atlas

Our analysis, with its unprecedented resolution, provides an invaluable framework for interpreting single-cell data from the *Drosophila melanogaster* central brain, enabling the exploration of relationships between developmental origins, timing, and functional identities (Figure 7A). To enhance accessibility and usability, we have deployed a user-friendly web portal (https://www.flycns.com), offering interactive web-based visualisation of the atlases generated in this study (Figure 7B). The portal includes broad atlases, such as those covering the whole head, whole head neurons, and central brain neurons. Additionally, specific sub-atlases are available for distinct neuronal populations, including adult type II neurons, early-born and late-born neurons, neuroendocrine cells, and monoaminergic neurons. More specialised sub- clustering atlases, such as those focused on individual hemilineages, are also available. Each atlas includes both known and predicted cell type annotations. Importantly, this intuitive interface enables easy visualisation of these single-cell transcriptomic data for the broad research community to explore cellular diversity and transcriptional complexity within the *Drosophila* central brain (Figure 7B). This study acts as an essential companion to the EM-resolution maps of the *Drosophila* central brain^10,11^, as integrating neuronal connectivity with molecular identity will enable a mechanistic understanding of how circuit structure links to function and ultimately behaviour.

**Figure 7.**
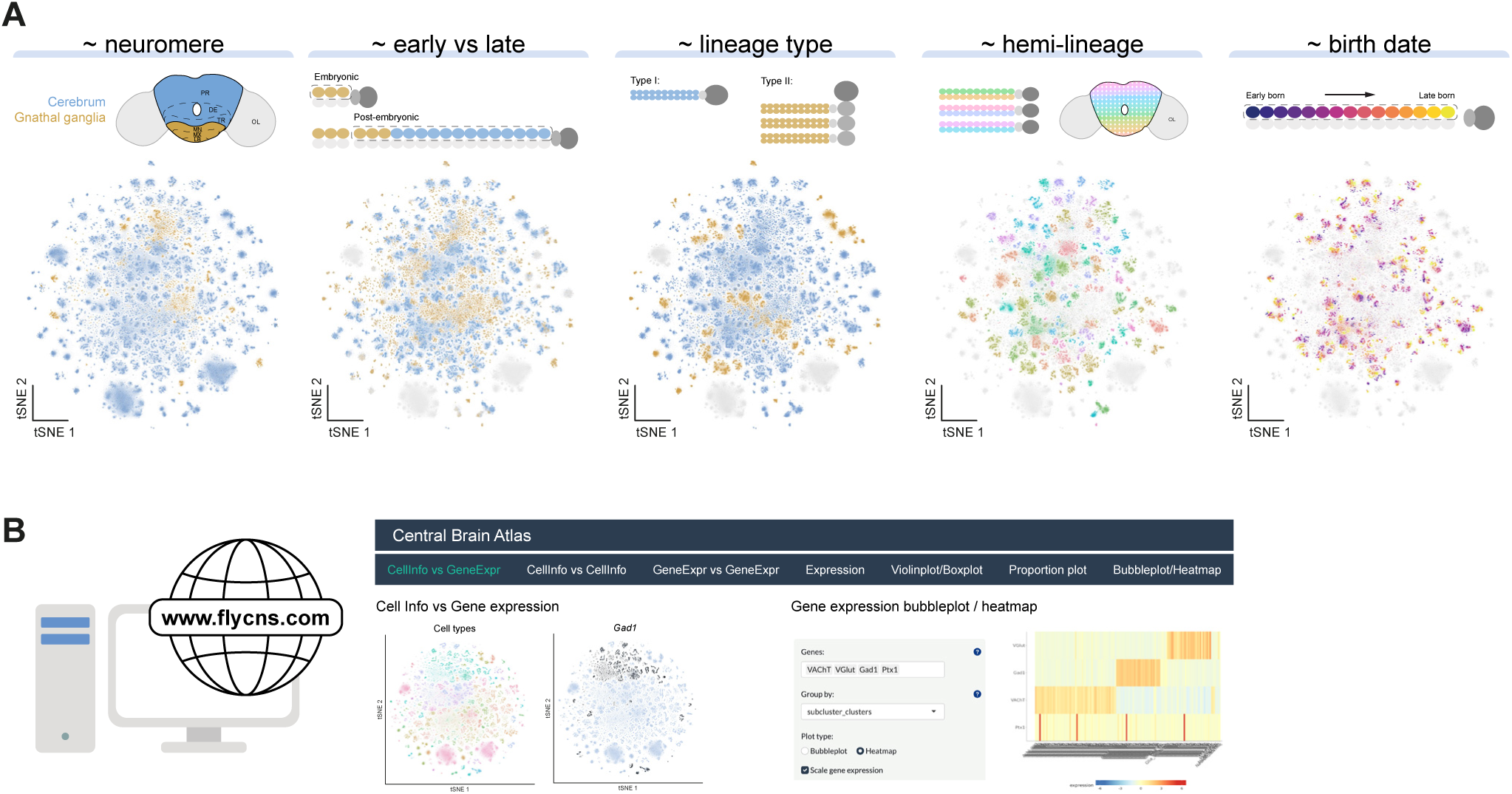
Interactive analysis of central brain neurons in *Drosophila*. (A) Organisational principles of the central brain neuron atlas. t-SNE visualisations of the central brain neuron meta-atlas classified across multiple analyses: neuromere, early vs. late birth, neuroblast type, hemilineage, and relative birth date within hemilineage. Schematics above each t-SNE illustrate the corresponding analysis. (B) Overview of the interactive web-based platform for exploring the central brain neuron atlas. Our web tool, available at flycns.com, provides an interactive interface for data exploration. Shown are examples of the interface: (left) the "Cell Info vs. Gene Expression" tab displaying two comparative t-SNEs annotated by subcluster and broad cell type categories; and (right) the "Gene Expression Bubbleplot/Heatmap" tab, which generates a heatmap illustrating the expression of multiple genes (*VAChT*, *VGlut*, *Gad1*, and *Ptx1*) across neuronal cell types in the central brain.

## DISCUSSION

Single-cell transcriptomic profiling technologies have revolutionised reductionist approaches to studying complex biological systems. Constructing comprehensive transcriptional atlases for complex organisms like *Drosophila melanogaster* is now an attainable goal, and the *Drosophila* research community has made substantial progress in this endeavour – most notably through the Fly Cell Atlas (FCA)^21^. Nevertheless, the diversity and complexity of cell types vary significantly across tissues, presenting unique challenges. The central brain remains one of the most complex tissues due to its vast array of highly specialised neuronal cell types, leading to its underrepresentation in current resources such as the FCA. We address this gap by generating a high-resolution single-cell transcriptional atlas of the *Drosophila melanogaster* central brain, identifying 4,167 distinct neuronal subtypes derived from 246 broader types (Figure S6). Integrating newly generated and existing single-cell datasets offers a scalable and cost-effective strategy for optimising available resources. This work represents a significant step forward in capturing the cellular diversity of the *Drosophila melanogaster* central brain, providing an invaluable resource for the research community.

Our unbiased investigation into the transcriptional characteristics defining neuronal cell types in the central brain revealed that shared developmental histories overwhelmingly emerge as the dominant feature. Transcriptionally defined cell types largely correspond to neuroblast-derived hemilineages (Figure 2). During development, these lineages are characterised by unique combinations of transcription factor expression, some of which remain throughout adulthood, giving each lineage a unique "genetic address"^63^. Our findings in the brain mirror those from our previous study on the ventral nerve cord^24^. Many of our predictions regarding hemilineage in that study have also been recently validated^30^. Future studies will undoubtedly further validate the patterns seen and discussed here.

A further defining feature, only apparent in our high-resolution atlas, is the striking transcriptional distinctness between early-born and late-born neurons. Neuroblasts progress through a series of transcription factor expression windows that govern the diversity of neurons they generate. This transcriptional progression acts as a temporal recorder of neuroblast divisions, with *Imp* marking neurons from early stages and *dati* marking neurons from later stages. In our atlas, early-born (*Imp*+) neurons and late-born (*dati*+) neurons form transcriptionally distinct classes (Figure 3; Figure S7-S8), with early-born neurons exhibiting more complex and distinct transcriptional profiles compared to late-born neurons, which often share overlapping profiles within their hemilineages. Early-born neurons have been found to have uniquely elaborate projection patterns and are proposed to serve as pioneering neurons, critical for establishing larval or adult brain architecture^64,65^. In contrast, late-born neurons integrate into pre-existing circuits, increasing the diversity of cell types. Thus, differing transcriptional complexities may reflect these differing developmental demands. Furthermore, in the recent adult brain connectome, it was found that known embryonically born neurons have significantly larger cell bodies and cell body fibres and often were unable to reliably be assigned to a lineage given their often-unique morphologies, indicating that their unique identities remain in the adult^10,11^. We show that genetic intersections which capture entire hemilineages can be deduced from our single-cell atlas, thereby integrating anatomical structures with their developmental origins in the central brain.

As developmental timing appeared to shape transcriptional relationships among neurons in the adult brain, we examined genes expressed within predicted hemilineages across pseudotime. We found that many temporal transcription factors expressed during development are maintained in mature neurons, reinforcing a developmentally rooted temporal transcription factor code that underlies much of the cellular diversity in the central brain (Figure 4). For example, the Mamo > Pdm3 > Br-Z4 > Bab1/2 > Pdm3 temporal cascade identified in our analysis highlights how developmental mechanisms governing temporal transitions during neurogenesis ultimately shape hemilineage subtypes in the adult brain, as previously described in the embryo^14^. Therefore, transcription factor expression not only identifies transcriptionally unique cell types (Figure 2) but also allows for refined sub-clustering of hemilineages (Figure 4), expanding the resolution to thousands of distinct neuronal types (Figure S6) and reflecting the massive diversity of the central brain. Overall, our findings support the neuronal lineage temporal patterning model previously proposed^64^, where neuronal complexity decreases over time across hemilineages through a progressive code of broad transcription factor expression across early- to late-born neurons, while recurring transcription factors and a mix of additional transcription factors, combine to diversify temporal windows.

While developmental histories broadly define transcriptional identities, our atlas also identified neuronal cell types that exhibit transcriptional convergence due to shared physiological roles despite arising from diverse developmental origins. This difference is particularly evident in neuroendocrine and monoaminergic neurons, which form discrete transcriptional clusters shaped by their shared function rather than lineage history (Figure 6, Figure S13). Focusing on neuroendocrine cells – a small but functionally essential subset – we demonstrate the ability of our atlas to resolve ultra-rare populations, such as Hugin-expressing neurons, and link them to anatomical identities. These findings suggest that functional specialization, particularly in peptidergic systems, can override developmental history in defining transcriptional identity.

Given that much of the central brain remains unannotated, our atlas and analytical approaches lay the foundation for systematically identifying and classifying all transcriptionally distinct cell types within the central brain and linking them to their anatomical counterparts. Our dataset categorises neurons based on multiple genetically identifiable characteristics, including neuromere location, neuroblast origin, birth order, and physiological properties such as neurotransmitter expression. To date, the transcriptional identity of central brain lineages has remained poorly defined. This study provides a critical pipeline for the systematic annotation of hemilineages, allowing for a more complete understanding of the brain’s developmental architecture. Additionally, we present a pipeline for linking transcriptionally defined cell types to their anatomical identities via genetic intersections, which can then be mapped onto the connectome (Figure 2). Furthermore, we demonstrate that transcriptionally and anatomically defined cell types can now be reinterpreted in the context of the entire central brain (Figure 5), enabling detailed comparisons between neuronal cell types. Integrating transcriptional profiles with the connectome provides a powerful approach for predicting how neurons communicate based on the genes they express. These insights will feed directly into systems neuroscience models, offering valuable information about the brain’s operation.

This study defines core heuristics for classifying neuronal identity in the central brain, offering a blueprint for integrating transcriptional, anatomical, and functional modalities (Figure 7A). Current single-cell technologies face limitations in detecting more subtle, state-dependent changes in neuronal gene expression. However, the increasing resolution of these methods and the ability to integrate new datasets with our atlas holds promise for addressing these challenges in the future. Our findings underscore the importance of linking adult neuronal characteristics to their developmental origins. Future work exploring the developmental dynamics that drive central brain neuronal diversity will be crucial for understanding how transcriptional identity influences adult neuronal connectivity, physiology, and behaviour. Finally, our atlas also sets the stage for evolutionary comparisons, enabling cross-species analysis of neuronal lineages, molecular programs, and circuit architectures. Identifying conserved transcriptional programs, such as temporal transcription factor cascades, will be key to understanding how nervous systems evolve to meet species-specific ecological demands.

## STAR METHODS

### KEY RESOURCES TABLE

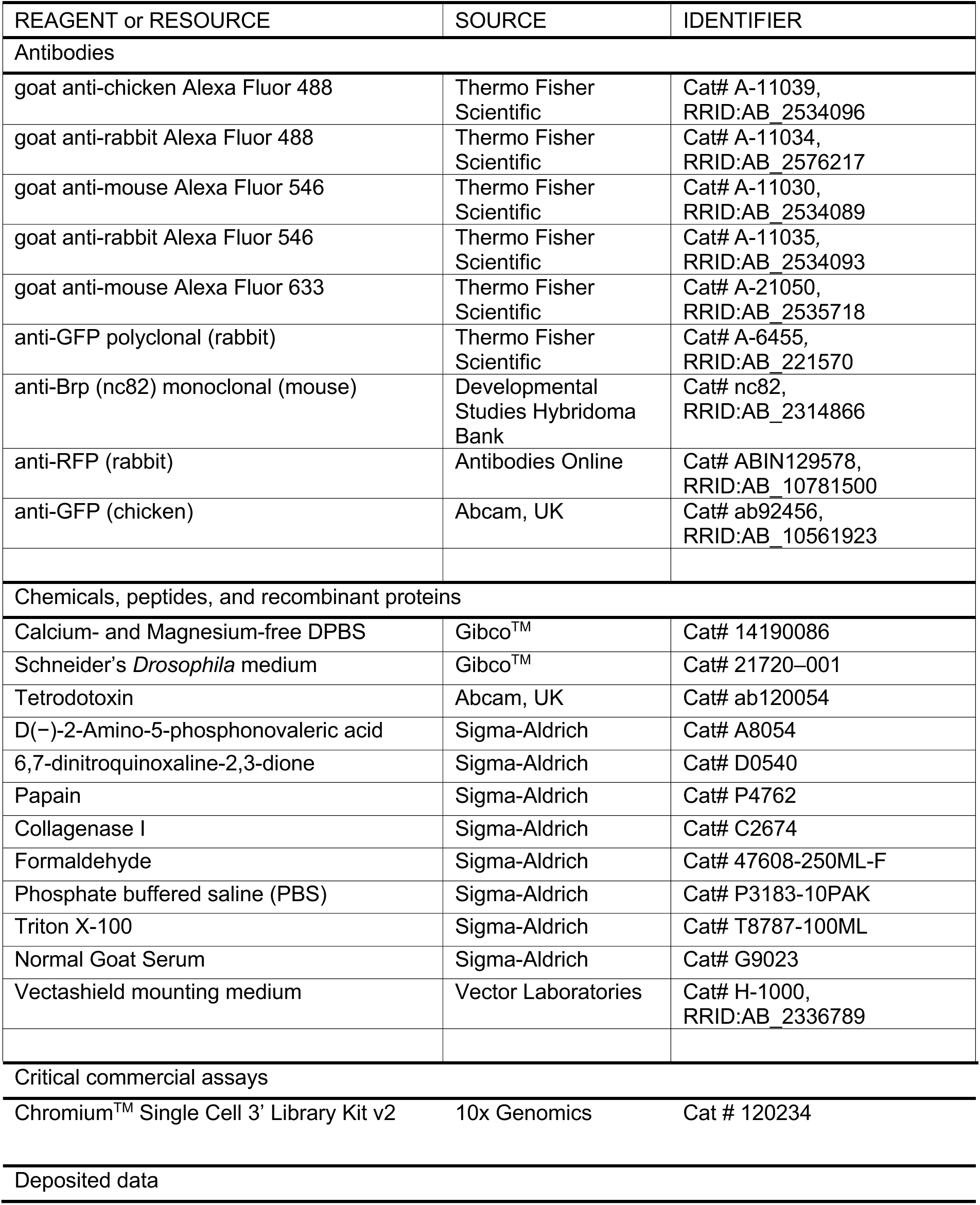
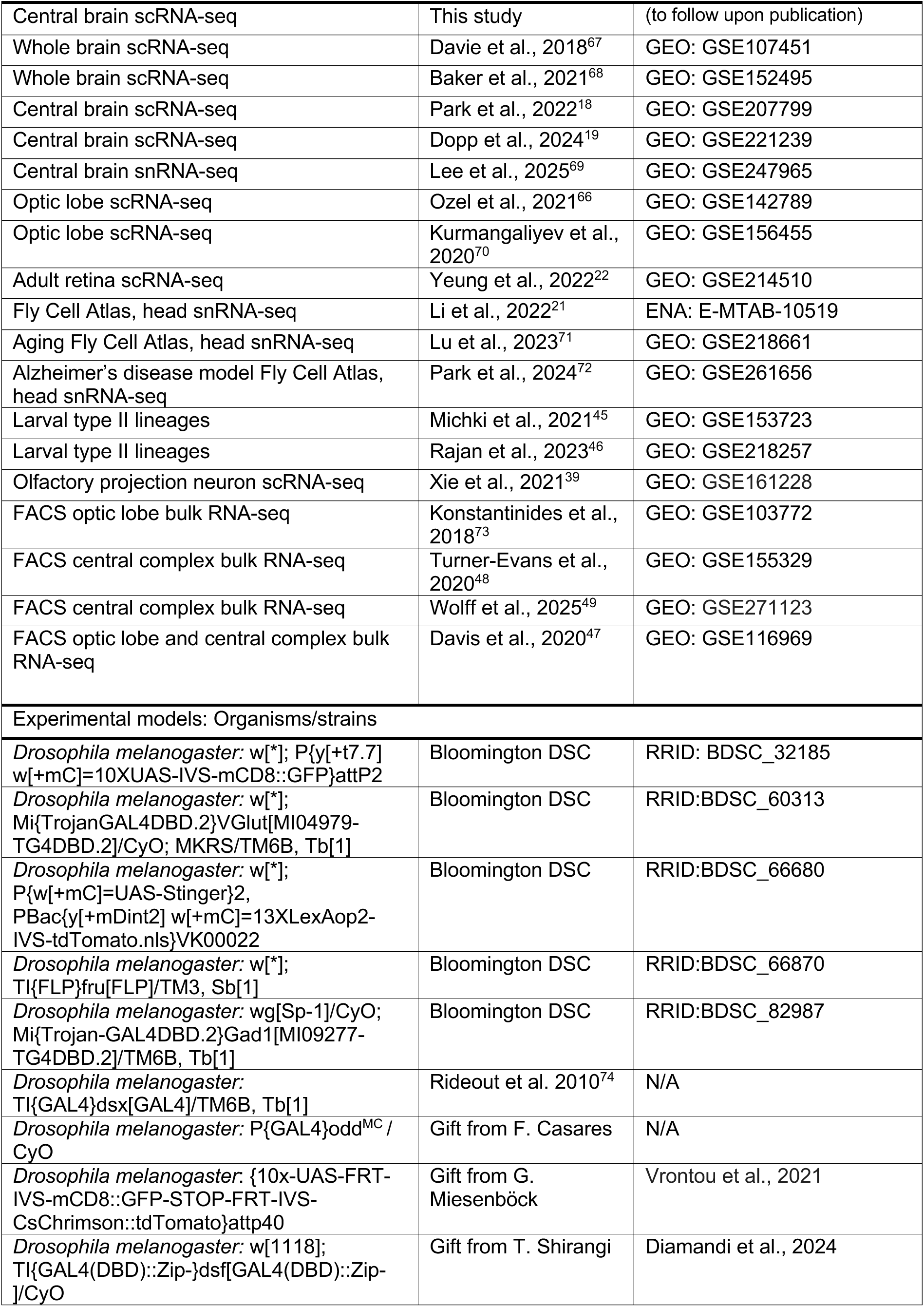
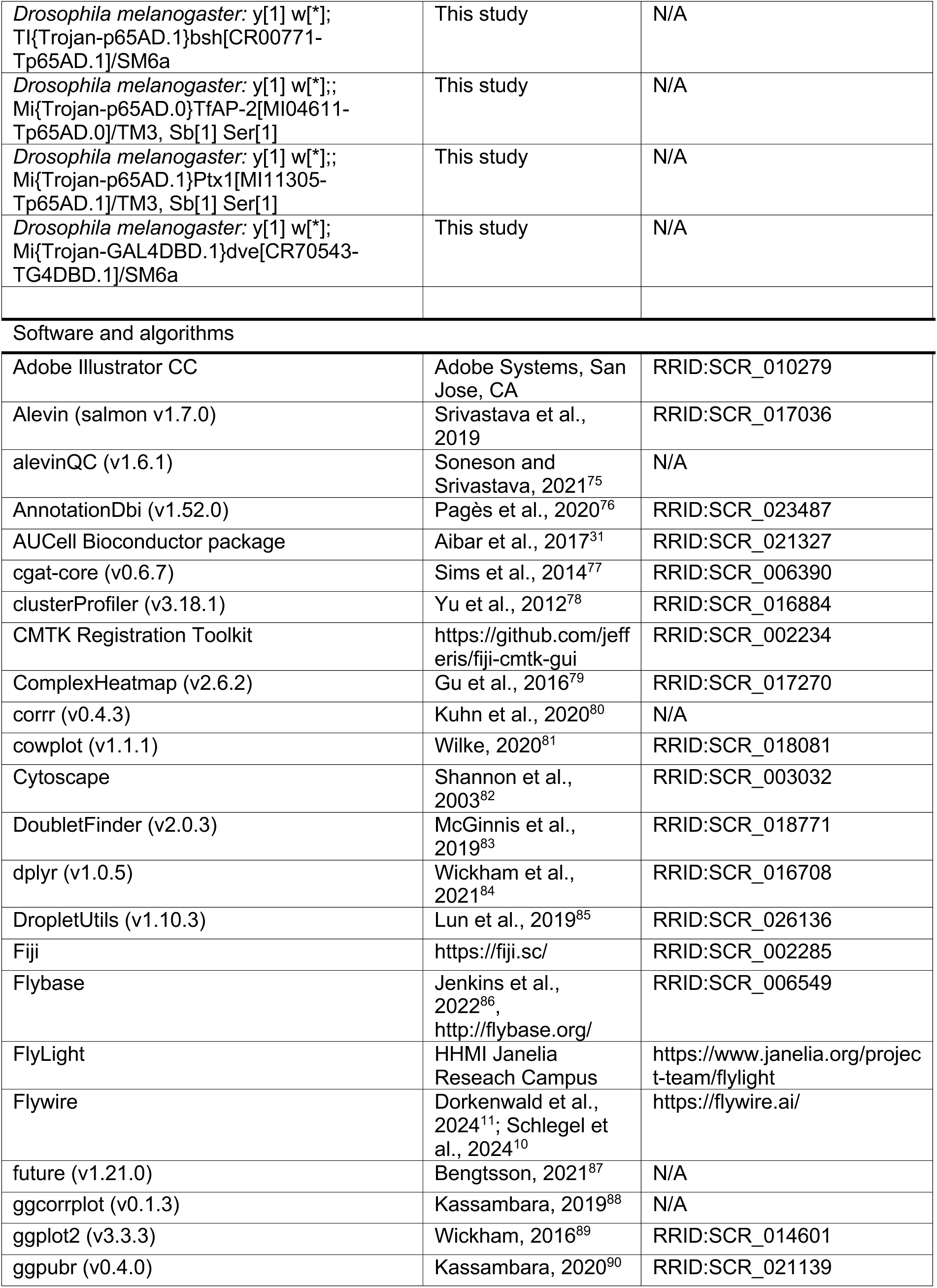
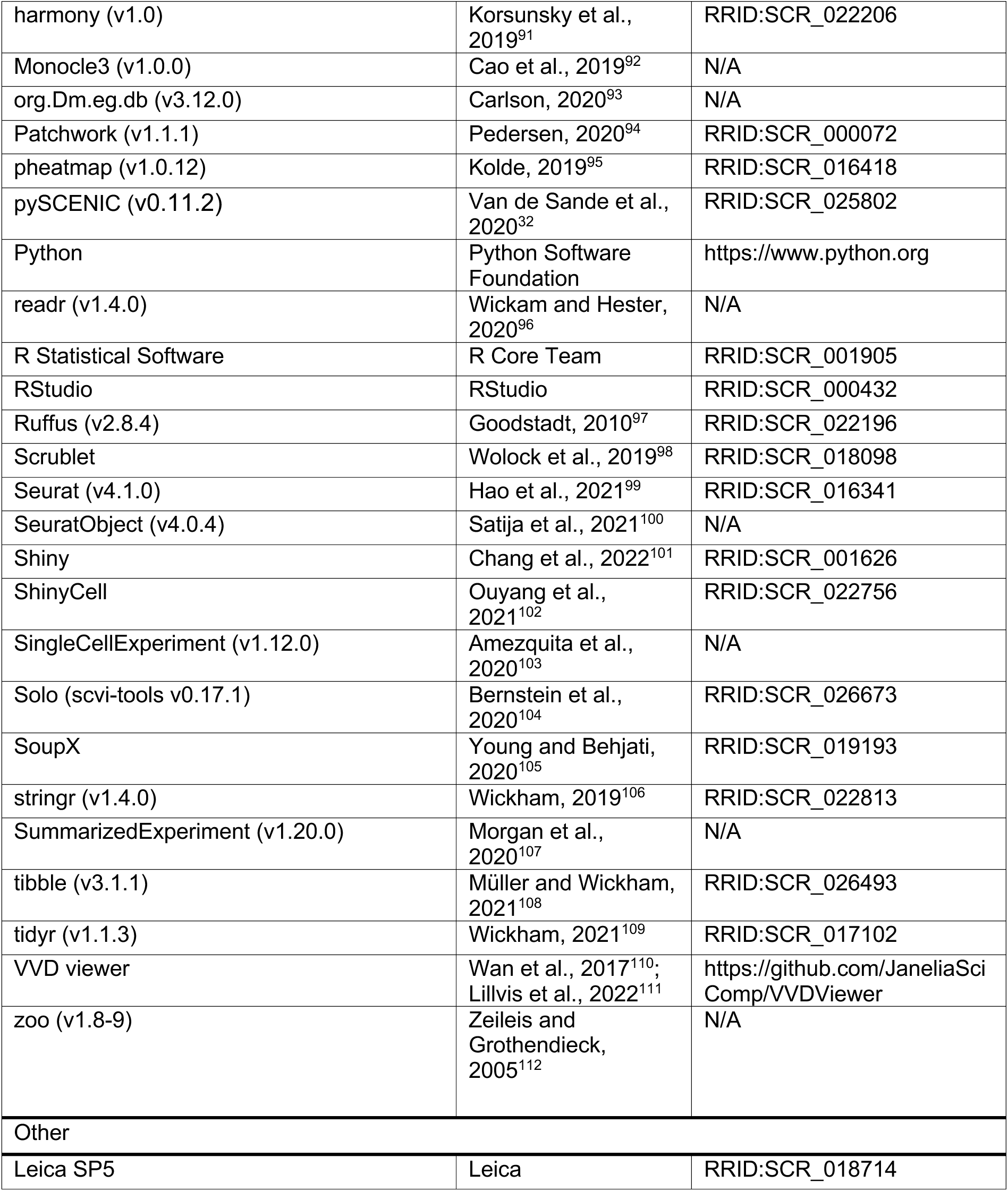

### RESOURCE AVAILABILITY

#### Lead contact

Further information and requests for resources and reagents should be directed to and will be fulfilled by the lead contacts Aaron M. Allen and Stephen F. Goodwin.

#### Materials availability

All unique/stable reagents generated in this study are available from the lead contacts without restriction.

### EXPERIMENTAL MODEL AND SUBJECT DETAILS

#### Drosophila stocks

All *Drosophila melanogaster* stocks were reared at 25°C and 40-50% humidity on standard cornmeal-agar food with a 12:12 light/dark cycle. Genotypes of the flies used were reported in the figure and legend. All strains used in the study are indicated in the key resources table.

#### Full Genotype List

**Related to Figures 1-2 and Figures S1-S4, SX.**

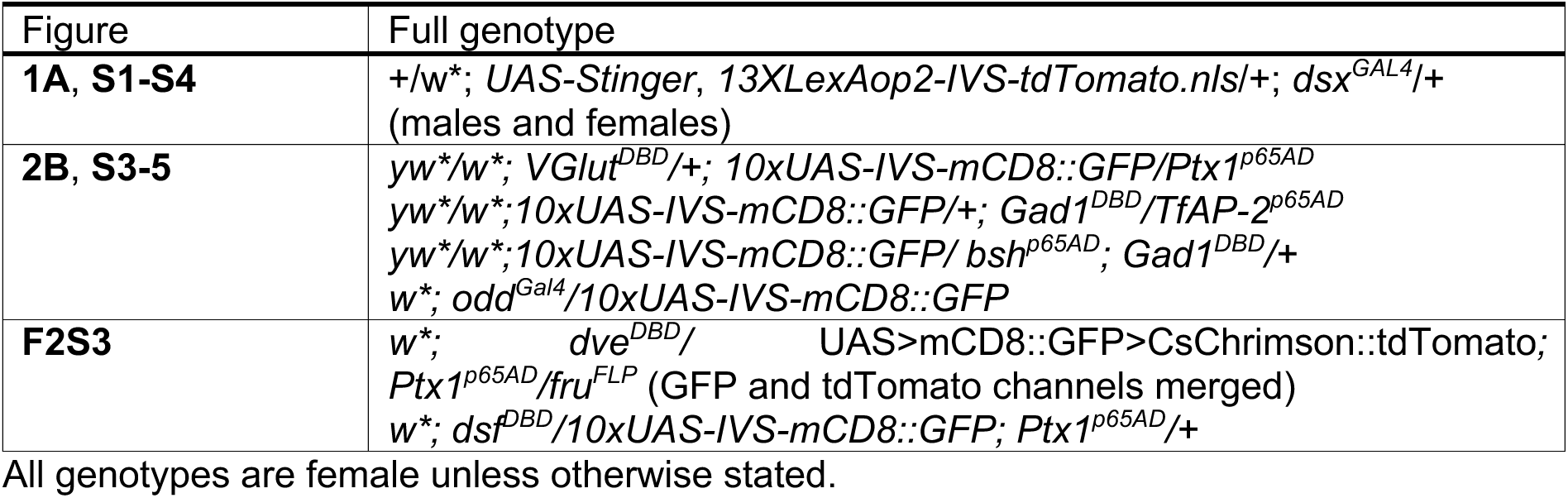

All genotypes are female unless otherwise stated.

### METHOD DETAILS

#### Central brain single-cell sample preparation

The central brain sample preparation was carried out as described previously^24^. The fly strain used *(+/w*; UAS-Stinger, 13XLexAop2-IVS tdTomato.nls/+; dsx^GAL4^/+*) was a genetic cross between *w*; UAS-Stinger, 13XLexAop2-IVS-tdTomato.nls* males (BDSC: 66680) and *+; +; dsx^GAL4^* virgin females^74^. Flies were raised at 25°C on standard cornmeal-agar food in an incubator with a 12:12 light/dark cycle. Virgin males and females were collected and stored individually. Flies were aged 5 days post-eclosion at 25°C prior to dissection. The central brain dissociation protocol was carried out as described previously (Allen et al., 2020). 2 replicates each of 20 male and 20 female central brains were individually dissected, removing optic lobes, in toxin- supplemented ice-cold calcium- and magnesium-free DPBS (Gibco^TM^, + 50 mM D(−)-2-Amino-5- phosphonovaleric acid, 20 mM 6,7-dinitroquinoxaline-2,3-dione and 0.1 mM tetrodotoxin). Each replicate was then washed in 1 mL ice-cold toxin-supplemented Schneider’s medium (tSM: Gibco^TM^ + toxins, as above). Brains were then incubated for 30 minutes in 0.5 mL of tSM containing 1 mg/mL papain (Sigma-Aldrich) and 1 mg/mL collagenase I (Sigma-Aldrich). Brains were washed once more with tSM and subsequently triturated with flame-rounded 200 mL pipette tips. Dissociated brains were resuspended into 1 mL PBS + 0.01% BSA and filtered through a 10 µm CellTrics strainer (Sysmex, 04-0042-2314).

#### Single-cell RNA-seq using 10x Chromium

Libraries were made using the Chromium Single Cell 3’ v2 kit from 10x Genomics. Cells were loaded in accordance with 10x Genomics documentation. The samples were sequenced with 8 lanes of Illumina HiSeq4000 by the Oxford Genomics Centre. For all code and specific details of what was run, please refer to https://github.com/aaron-allen/Dmel-adult-central-brain-atlas.

#### Alignment and Cell Identification

The fastq files were aligned to the r6.30 transcriptome release and included all transcript types. Digital expression matrices were generated using salmon-alevin tool for all the droplets per sample^113^ (v1.7.0). To identify cell-containing droplets, we used the "emptyDrops" function from the DropletUtils package^85^. This method models the ambient RNA profile from low-count barcodes (<100 UMI), assumed to represent empty droplets. Each droplet is then statistically tested for significant deviation from the ambient profile using a Monte Carlo simulation. Droplets with false discovery rate (FDR)-adjusted p-values below 0.1% were classified as containing real cells. Filtered cell-containing barcodes were written out into Matrix Market (.mtx) Cellranger v2 format using the "write10xCounts" for downstream processing in Seurat. Between 300,000,000 and 500,000,000 reads per sample were processed, with approximately 68-72% of reads mapping to the transcriptome and a mean deduplication rate of 58-69%. The results form alevin were inspected with alevinQC^75^ (v1.6.1).

#### Post-alignment Processing

Most downstream single-cell processing was performed in R (v4.0.2; R Core Team, 2021) with the Seurat package^99^ (v4.1.0), along with many other packages including: ComplexHeatmap^79^ (v2.6.2), corrr^80^ (v0.4.3), cowplot^81^ (v1.1.1), dplyr^84^ (v1.0.5), future ^87^ (v1.21.0), ggcorrplot^88^ (v0.1.3), ggplot2^89^ (v3.3.3), ggpubr^90^ (v0.4.0), Patchwork^94^ (v1.1.1), pheatmap^95^ (v1.0.12), readr^96^ (v1.4.0), SeuratObject^100^ (v4.0.4), Shiny^101^, SingleCellExperiment^103^ (v1.12.0), stringr^106^ (v1.4.0), SummarizedExperiment^107^ (v1.20.0), tibble^108^ (v3.1.1), tidyr^109^ (v1.1.3), zoo^112^ (v1.8-9). Droplets were initially filtered to have greater than 500 UMI, greater than 300 genes, less than 15% mitochondrial UMI, and less than 15% heat-shock UMI. Doublets were predicted using 3 different methods: DoubletFinder^83^ (v2.0.3), Scrublet^98^, and Solo^104^ (scvi-tools v0.17.1). Droplets were annotated as doublets if at least two of the three methods predicted them to be so (Figure S1D- E). All three methods showed similar behaviour in predicting doublets across distinct cell types (Figure S1E-F). We next estimated and removed ambient RNA contamination using SoupX^105^ (v1.5.0). Ambient estimation was performed using sets of marker genes for mutually exclusive cell types (Figure S1G-J). The data was log-normalised using "NormalizeData", and 1000 variable genes were selected with "FindVariableFeatures" using method "mean.var.plot". The normalised counts were scaled with "ScaleData", and a principal component analysis was performed using "RunPCA". The replicates were batch-corrected and integrated with Harmony (v1.0; Korsunksy et al., 2019) using "RunHarmony". Uniform Manifold Approximation and Projections (UMAPs) and t-distributed stochastic neighbour embeddings (t-SNEs) were computed with "RunUMAP" and "RunT-SNE". Unsupervised clustering was performed by running "FindNeighbors" and "FindClusters" using the Leiden algorithm.

#### Re-processing of publicly available scRNA-seq data

Fastq files were downloaded from NCBI’s Sequence Read Archive (SRA) and EMBL-EBI’s European Nucleotide Archive (ENA) for the following datasets^18,19,21,22,66–72^. These datasets were all 10x Chromium 3’ scRNA-seq chemistries but varied in which version (v2 and v3), as well as in tissue type (dissected whole brain, dissected central brain, dissected optic lobe, dissected retina, and whole head), and cellular preparation type (whole cell and nuclei). Although our primary focus was to achieve a central brain neuronal atlas, we included datasets of dissected and enriched optic lobe and retina samples to aid in the identification of these cell types in the central brain dissected and whole head preparations. Both the Ozel et al. (2021)^66^ and Kurmangaliyev et al. (2020)^70^ included dissected tissue throughout pupal development, but we only used the pharate adult and adult time points here. All datasets were re-processed as above to be consistent with our dataset and for ease of integration (Figure S2A).

#### Integrating Datasets

To integrate these datasets, we annotated and removed doublets but decided not to ambient correct the data. When estimating ambient contamination with SoupX, as detailed above (Figure S1), some of the datasets had predicted ambient contamination over 70%, particularly those in Park et al. (2024)^72^. Removing the estimated ambient contamination of these high ambient samples resulted in most cells being removed by UMI and gene filtering. And so, we opted not to correct for ambient contamination and instead to be vigilant in downstream analyses, ensuring that gene expression profiles were consistent across datasets. All the individual samples from each of the datasets were integrated with Harmony, using "RunHarmony" with the inclusion of dataset and preparation type (cell vs. nuclei) metadata to be corrected for, along with individual sample ID. Once integrated into a meta-head atlas (Figure S2B), non-neuronal cell types were annotated by marker gene expression and the proportion of each cluster originating from each tissue type (Figure S2C). These neuronal cells were then extracted, re-integrated, and re- clustered, generating a meta-neuronal atlas (Figure S2D). As before, peripheral and optic lobe- specific cell types were annotated by marker gene expression and cluster-wise proportion of tissue type (Figure S2E), as well as correlating to bulk-RNA-seq of known specific cell types (data not shown) and transferring the existing annotations from Janssens et al., 2022 (which was a re- analysis of the data from Davie et al., 2018) and Ozel et al., 2021 (Figure S2F). With the peripheral and optic lobe neurons annotated, the remaining central brain neurons were extracted, re- integrated, and re-clustered, generating a meta-central brain neuronal atlas of 329,466 cells/nuclei (Figure 1A; Figure S2G). A range of cluster resolutions was explored (data not shown), and a resolution of 10 was chosen for the next steps.

#### Annotation of sex

Two of the datasets used in our meta-analysis did not separate the sexes into separate samples^18,67^. As a result, we’ve had to rely on in-silico sexing of these samples. We used Seurat’s "AddModuleScore" function with the male-specific genes *lncRNA:roX1* and *lncRNA:roX2*. Cells with a module score less than zero were labelled female, and cells with module scores greater than zero were labelled male. To test the efficacy of this method, we performed *in silico* sexing of the datasets where the sexes were processed separately and achieved a precision of 0.93-1.00 and a recall of 0.89-0.97 for identifying a cell as female, depending on the dataset.

#### Annotation of broad cell types

Broad neurotransmitter identity (cholinergic, glutamatergic, GABAergic, monoaminergic, and neuroendocrine) and special case cell types (Kenyon cells and motor neurons) were annotated using sets of marker genes. Seurat’s "AddModuleScore" function was used with sets of genes specific to each broad cell type to calculate the average expression level of these gene programs across all cells. These module scores were then l2-normalised. Cells were annotated as a given broad cell type by both having a positive module score for that gene program and that l2- normalised module score being higher than any other score. The most common annotation for each cluster was then extrapolated to the remaining cells in each cluster at resolution 10. Each annotated broad cell type was then sub-clustered by extracting, re-integrating, and re-clustering only those cells (data not shown). These new sub-clustered identities were then mapped back onto the main meta-central brain neuronal atlas.

#### Gene ontology analysis

There was a total of 16,267 genes of the 16,841 total genes that had at least one UMI detected in this meta dataset. To estimate the universe or background of truly expressed genes from those arising from contamination, additional filtering was imposed. To be classified as "expressed", a gene had to have the following: (1) Summed UMI across all cells greater than or equal to 400, (2) Maximum UMI observed within a cell greater than or equal to 4, (3) The percent contribution of UMI from each dataset less than or equal to 60%, and (4) The percent contribution of UMI from the Park et al. (2024)^72^ dataset less than or equal to 30%. This resulted in 9,017 "expressed" genes in the meta dataset. Significantly enriched genes in the clustered cell types were calculated with Seurat’s "FindAllMarkers" using the Wilcoxon Signed-Rank test and with an average log fold change greater than 0.5 and Bonferroni-adjusted p-value less than 0.05. Gene ontology analysis was performed using the packages "clusterProfiler"^78^ with "org.Dm.eg.db"^93^ and " AnnotationDbi"^76^. The "enrichGO" function was used to determine the enriched terms for each of the molecular function, biological process, and cellular compartment categories (Figure S3F).

#### Gene regulatory network analysis with SCENIC

pySCENIC^32^ (v0.11.2) was run on select, individual subclustered clusters (Figure 2C; Figure 6) to identify transcription factor (TF) based regulons by using the gene expression data as input. Briefly, the raw expression matrix was filtered to retain genes expressed in >1% of cells and with a count >3 × 0.01 × number of cells. Modules comprising transcription factors and co-expressed genes were generated using GRNBoost2, then pruned to remove indirect targets lacking enrichment for the corresponding transcription factor motif and further refined (cisTarget). To build the final set of TF regulons, the predicted target genes of each TF module that show enrichment of any motif of the given TF are then merged. Due to stochasticity in gene regulatory network inference using GRNBoost2, each pySCENIC run can identify a different number of regulons and different target genes for each transcription factor. Thus, pySCENIC was run 100 times. High-confidence regulons were defined as those that occurred in >50% of runs and contained at least five high-confidence target genes. High-confidence target genes were those found within a regulon in >50% of runs. Cells were then scored for the activity of each high-confidence regulon (including only high-confidence target genes) using the AUCell Bioconductor package^31^. Briefly, single cells are scored by calculating the enrichment of a regulon, which is measured as the Area Under the recovery Curve (AUC) across the ranking of all genes in a particular cell, whereby genes are ranked by their expression values. AUC represents the proportion of expressed genes in the signature and their relative expression values compared to the other genes within the cell. The output of this step is a matrix with the AUC score for each gene set in each cell. We used either the AUC scores (across regulons) directly as continuous values to cluster single cells or a binary matrix generated using a cutoff of the AUC score for each regulon. These steps were run as a pipeline written in the cgat-core ruffus framework^77,97^. As described above, AUC scores per cell were fed into Seurat and processed as a separate assay. Cell-type enriched regulons were calculated using Seurat’s "FindAllMarkers" function. Gene regulatory networks were visualized with Cytoscape^82^. For visualization purposes, GRNs were further filtered to only include positive regulons less than 100 genes in size, and each node (gene) had to have a maximum expression of at least 3 UMI and total expression of at least 80 UMI.

#### Early- vs late-born morphology quantification

To annotate early-born ("*Imp*>*dati*") and late-born ("*Imp*<*dati*"), we used Seurat’s "AddModuleScore" to calculate the average expression level of early (*Imp*, *mamo*) and late (*dati*, *pros*) gene programs across all cells. A cell was annotated as early-born if the l2-normalised early- born module score was both greater than zero and greater than the l2-normalised late-born module score. Conversely, a cell was annotated as late-born if the l2-normalised late-born module score was both greater than zero and greater than the l2-normalised early-born module score. We used three separate methods to quantify the "punctate" vs "serpentine" t-SNE morphology differences between early-born and late-born neurons: (1) Mean pairwise distance, (2) Fragmentation score, (3) Modularity score (Figure 3C-D, Figure S8F-I). Before calculating these metrics, Kenyon cells and cells where the early- and late-born module scores were either both negative or equal were removed. The mean pairwise distance in t-SNE space between all cells within each cluster was calculated with the "dist" function. Cells were then split into early-born ("*Imp*>*dati*") and late-born ("*Imp*<*dati*"), as described above (Figure 3C). Secondly, the fragmentation score is the number of subclusters generated using the "dbscan" function with an epsilon neighbourhood radius ("eps") of 0.5 and the number of neighbourhood members ("minPts") equal to 5 (Figure 3D). A range of values for "eps" and "minPts" were tested, and they all generated a difference (Figure S8G-H). Thirdly, we calculated a modularity score for each cell in a cluster-wise fashion. Distances between points in t-SNE space were calculated with the "dist" function. Graphs were calculated with "graph_from_adjacency_matrix" and communities were detected with "cluster_fast_greedy". Lastly, the modularity of these communities was calculated with the "modularity" function (Figure S8I). The differences were statistically quantified using the Wilcoxon Signed-Rank test with Bonferroni correction.

#### Annotating and subclustering ALad1

The ALad1 olfactory projection neuron lineage is co-positive for *acj6* and *Oaz*^21^. We extracted the cells from cell type "Achl 032", the only cluster co-positive for these genes, as an initial proxy. These data were integrated, as above, with previously published scRNA-seq data from the developing olfactory projection neurons from multiple hemilineages^39^ that were downloaded from the Gene Expression Omnibus (GSE161228). In this integration of datasets, the hemilineage identity, annotated in Xie et al. (2021)^39^, was extrapolated to the cells from the other datasets by cluster. Clusters from the meta-central brain neuron atlas now annotated as ALad1 (*acj6+*, *Oaz+*, and *vvl-*) were extracted, re-integrated, and re-clustered, as above, to form our refined proxy for the adult ALad1 hemilineage (Figure S9).

#### Pseudotime analysis

Pseudotime analysis was performed using Monocle3^92^ (v1.0.0) using standard methods. Briefly, the data were processed with functions "new_cell_data_set", "preprocess_cds", "align_cds", "reduce_dimension", "cluster_cells", and "learn_graph" using default settings. Cells were ordered with "order_cells" setting all *Imp*+ cells as the root. Genes with significantly variable expression across pseudotime were calculated with "graph_test" and filtered for q-value less than 1e-50 and Moran’s I greater than 0.2. Heatmaps of significantly varying genes were plotted with the package "ComplexHeatmap". For the systematic pseudotime analysis across the central brain cell types (Figure 4H-J), only clusters at Leiden resolution 10 that met the following criteria were used: (1) predicted type I or type II lineages, (2) l2-normalised late-born module score (see above) greater than zero and greater than l2-normalised early-born module score, (3) a single sub-region of the cluster positive for *br*. A total of 84 of the 220 clusters met these criteria and were subclustered and subjected to pseudotime analyses. Following the standard processing described above, the resulting pseudotime metadata for each of these clusters was normalised to a percent to be comparable to each other. To identify genes that repeatedly temporal transcription factors that vary over pseudotime, we filtered the resulting pseudotime markers for each of the clusters for transcription factors that varied significantly in at least 25 of the 84 clusters (Figure 4J).

#### Annotating and subclustering type II NSC lineages

To annotate neurons derived from type II lineages in our central brain neuronal atlas, we leveraged previously published scRNA-seq of FAC-sorted late L3 larval type II lineages^45,46^. Fastq files from these datasets were downloaded and re-processed as described above. After an initial integration, the central brain neuronal population of these data were extracted, re-integrated, and re-clustered to form a larval type II neuronal atlas. These larval data were then integrated with a subset of our re-processed adult datasets. For this integration, we selected only whole-cell preparation datasets with the Kenyon cells removed (as they are derived from their own type of neuronal stem cell). Resulting clusters at resolutions of 0.2, 1, 2, 4, and 10 that have more than 5% of their cells derived from the larval type II datasets were annotated as "type II", and all others were annotated as "type I" (Figure 5B). These lineage-type annotations were then extrapolated, in a cluster-wise fashion, to the rest of the meta-central brain neuron atlas (Figure 5C). Once annotated, these annotated type II neurons were extracted, re-integrated, and re-clustered, as above, to generate an adult type II neuronal stem cell lineage atlas (Figure 5D). Type II cell types were annotated by a combination of correlation to bulk-RNA-seq data (as above, Figure S10), marker gene expression, and genetic intersection^20^ (Figure 2).

#### Annotating and subclustering neuroendocrine and monoaminergic neurons

Neuroendocrine clusters were annotated as clusters with a positive z-scored expression of the transcription factor *dimm*. These cells were extracted, re-integrated, and re-clustered as described above. After this initial subclustering, a few cell types were negative for *dimm* as they were part of a heterogeneous parental cluster. These cells were removed, and the remaining cells were re-subclustered. Gene regulatory network analysis was performed on the neuroendocrine cell types as described above. Subclustering of the monoaminergic neurons was conducted in a similar fashion. Broad monoaminergic class was assigned by the following marker genes: dopaminergic (*ple*+ and *DAT*+), histaminergic (*Hdc*+), octopaminergic (*Tdc2*+ and *Tbh*+), serotonergic (*SerT*+ and *Trh*+), and tyraminergic (*Tdc2*+ and *Tbh*-). Further subclustering of the non-PAM (Figure S13F-G) and PAM (Figure S13H-J) subdivision of monoaminergic clustering was performed by separating the non-PAM cells from the PAM (*DAT*+, *Fer2*+, and *Imp*-) cells.

#### Generating a web app

Our interactive web tool for visualization of these scRNA-seq data was generated using a modified version of "ShinyCell", an R-based Shiny App^102^; https://github.com/SGDDNB/ShinyCell). Specifically, we forked and modified "easyshiny" (https://github.com/NBISweden/easyshiny), which is a forked version of the original "ShinyCell".

#### Generation of split-Gal4 driver lines

We used existing coding intronic Minos-mediated integration cassette/CRISPR-mediated integration cassette (MiMIC/CRIMIC) lines^114,115^ (see Key Resources Table) to generate split- GAL4 drivers using the Trojan method^116^. pBS-KS-attB2-SA()-T2A-Gal4DBD-Hsp70 or pBS-KS- attB2-SA()-T2A-p65AD-Hsp70 vectors^116^ with the appropriate reading frame were inserted in the MiMIC/CRIMIC locus of a given line via recombinase-mediated-cassette-exchange through injection (BestGene, CA). Stocks were generated with the transformed flies as described in^116^.

#### Immunohistochemistry

After a brief pre-wash of adult flies in 100% EtOH to remove hydrophobic cuticular chemical compounds, brains were dissected in PBS at RT (20-25°C), collected in 2 mL sample tubes and fixed with 4% formaldehyde (Sigma-Aldrich) in PBS (Sigma-Aldrich) for 20 min at RT. After fixation, tissues were washed in 0.5-0.7% Triton X-100 in PBS (Sigma-Aldrich) (PBT) 3 times each for 20 min at RT. After blocking in 10% normal goat serum (Sigma-Aldrich) in PBT (NGS/PBT) overnight (8-12 h) at RT, tissues were incubated in primary antibody solutions for 48-72 hrs at 4°C (1:1000, rabbit anti-GFP, Thermo Fisher Scientific; 1:1000, chicken anti-GFP, Abcam; 1:1000, rabbit anti- RFP, antibodies-online; 1:10, mouse anti-Brp, Developmental Studies Hybridoma Bank; 1:500). After 4 washes in PBT for 1 hr each at RT, tissues were incubated in secondary antibody solutions for 48 h at 4°C (1:500, anti-rabbit Alexa Fluor 488, anti-chicken Alexa Fluor 488, anti-mouse Alexa Fluor 546, anti-rabbit Alexa Fluor 546, anti-mouse Alexa Fluor 633, Thermo Fisher Scientific). After 4 washes in PBT for 1 hr each at RT specimens were imaged directly or 70% glycerol in PBS was added to the sample tubes, which were subsequently transferred to -20°C and kept for at least 8 hr for tissue clearing. Specimens were mounted in Vectashield (Vector Laboratories).

#### Confocal image acquisition and processing

Confocal image stacks were acquired on a Leica TCS SP5 confocal microscope at 1024 x 1024- pixel resolution with a slice size of 0.29 µm or 1 µm. Water-immersion 25x and oil-immersion 40x objective lenses were used for brain images. Images were registered onto the intersex template brain using the Fiji Computational Morphometry Toolkit (CMTK) Registration GUI (https://github.com/jefferis/fiji-cmtk-gui). For segmented images we used the software VVD Viewer^110,111^ (https://github.com/JaneliaSciComp/VVDViewer) to render the registered image stacks in 3D, manually mask other neurons co-labelled in the image and segment out neurons of interest. All segmented images include unsegmented versions within the supplemental figures.

## DATA and CODE AVAILABILITY

The datasets generated and described here can be visualised and explored at https://www.flycns.com/. Raw sequencing files (fastq) and digital expression matrices from each replicate will be available from the Gene Expression Omnibus. The code used in this analysis will be made available upon publication. The following data sets were generated: (to follow upon publication). The following previously published data sets were used: GSE107451, GSE152495, GSE207799, GSE221239, GSE247965, GSE142789, GSE156455, GSE214510, E-MTAB-10519, GSE218661, GSE261656, GSE153723, GSE218257, GSE161228, GSE103772, GSE155329, GSE271123, GSE116969. Any additional information needed to reanalyse the data reported in this paper is available from the lead contact by request.

## ACKNOWLEDGEMENTS

We thank D. Shepherd, S. Russell, E. Rideout, E. J. Clowney, Y. Ding, J. Walsh, and the Goodwin Lab for helpful discussions and critical reading of the manuscript. We are grateful to F. Casares, T. Shirangi, and G. Miesenböck for sharing important stocks and reagents with us. We would like to thank S. Liu for early access to the data from^19^. Stocks obtained from the Bloomington Drosophila Stock Center (NIH P40OD018537) were used in this study. We thank A. Rings, S. Birtles, V. Croset, and C. Treiber for dissection/dissociation assistance. Sequencing was carried out by The Oxford Genomics Centre. A.M.A. and M.C.N. were supported by a Wellcome Trust Senior Investigator Award to S.F.G (106189/Z/14/Z) and a grant from the BBSRC (BB/X016595/1) to M.C.N., A.M.A., and S.F.G. T.N., and F.A. were supported by a grant from the BBSRC (BB/T001348/1) to T.N., and S.F.G. A.M.A., and D.A. were supported by a Wellcome Collaborative Award to D.S., and S.F.G. (209235/Z/17/Z).

## AUTHOR CONTRIBUTIONS

Conceptualization, A.M.A., M.C.N., and S.F.G.; Methodology, A.M.A., M.C.N., T.N., F.A., D.A., D.S., and S.F.G.; Investigation, A.M.A., M.C.N., T.N., D.A., and F.A.; Resources, S.F.G.; Writing, A.M.A., M.C.N., T.N., F.A., and S.F.G.; Supervision, D.S., and S.F.G.; Funding Acquisition, D.S., and S.F.G.

## DECLARATION OF INTERESTS

The authors declare no competing interests.

---

**Figure S1.**
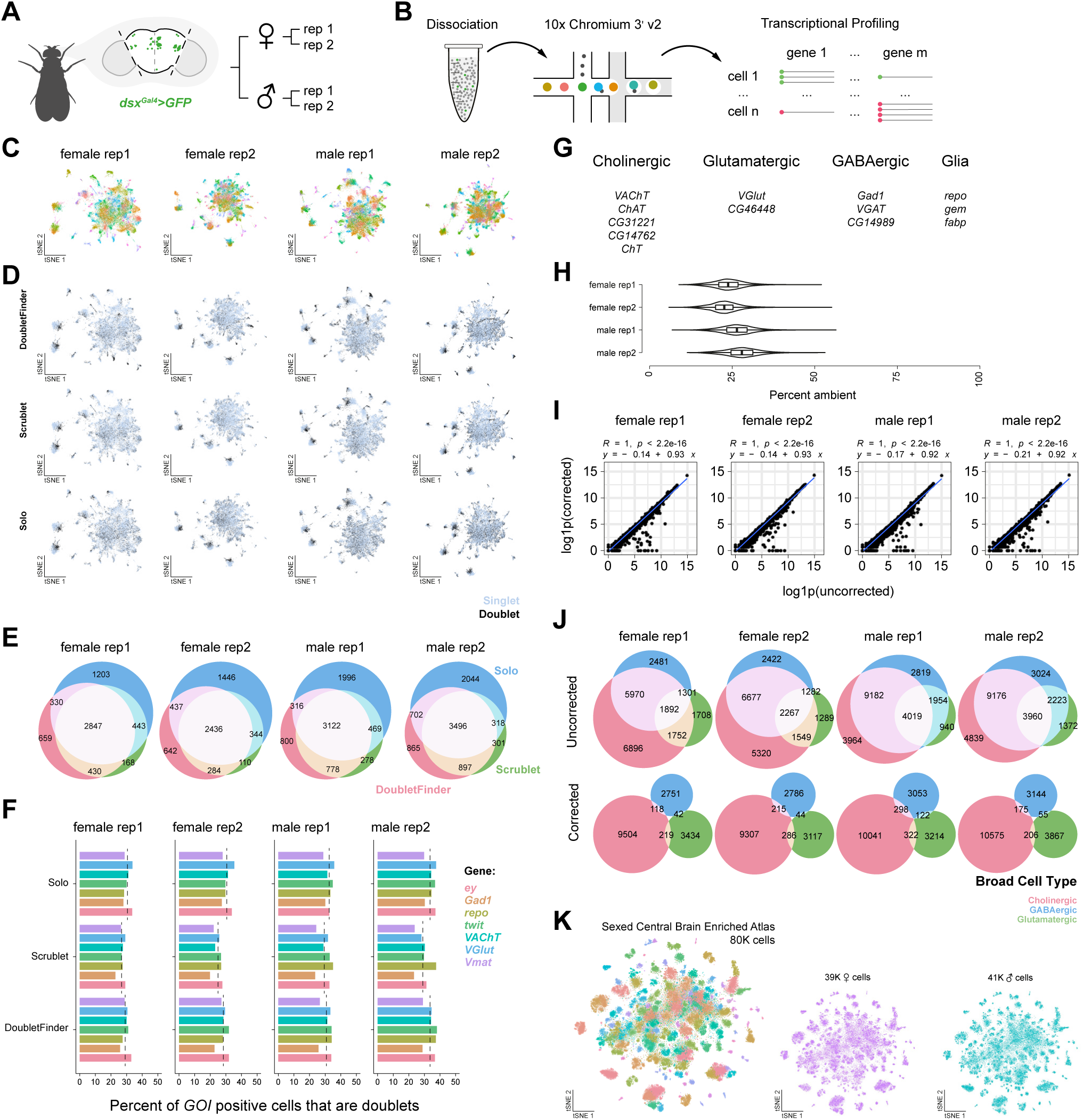
Generation of sexed central brains scRNA-seq dataset. Related to Figure 1. (A) Generation of sexed central brain scRNA-seq atlas: central brains expressing *doublesex* driven EGFP were dissected, dissociated, and analysed. (B) 10x Chromium 3’ v2 workflow for sexed single-cell atlas. (C) t-SNEs of individual replicate with cell types defined by unique colours. (D) Singlets (light blue) vs doublets (black) across replicates comparing alternative identification methods (DoubletFinder, Scrublet, and Solo). (E) Euler diagrams comparing doublets identified between methods per replicate. (F) Percent of GOI positive cells that are doublets by method, per replicate. (G) Sets of cell type-specific, mutually exclusive marker genes for broad cell types used for estimating ambient contamination. (H) Combination violin boxplots of the estimated percent ambient contamination in each replicate. (I) Per replicate correlation of pseudo-bulked uncorrected and corrected gene counts (UMI). (J) Euler diagrams comparing the co-expression of mutually exclusive markers for broad cell types with uncorrected (top) and corrected (bottom) gene expression profiles between replicates. (K) t-SNEs of sexed central brains atlas with cell types defined by unique colours (left) and by sex (right).

**Figure S2.**
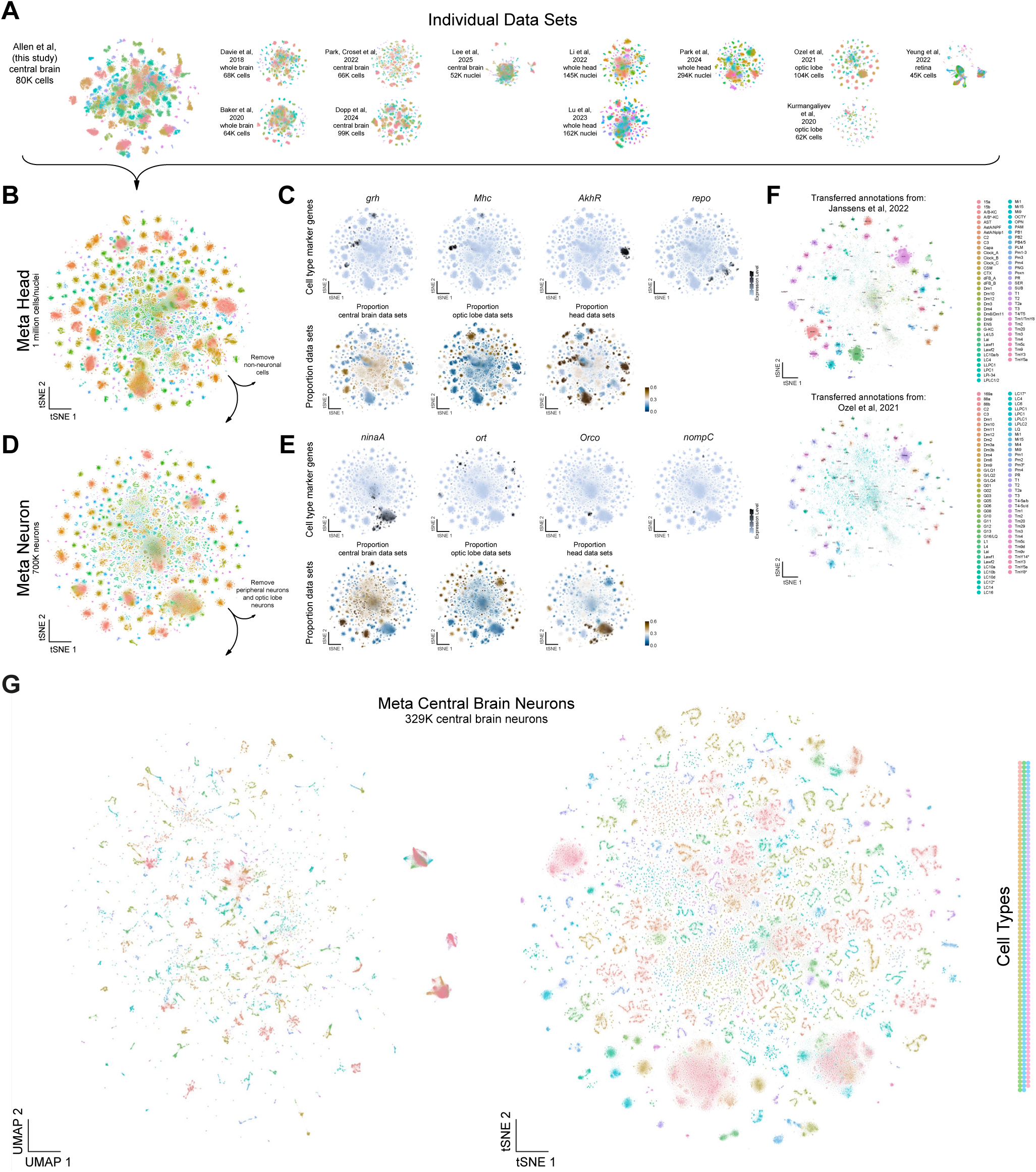
Integration and cell type identification within the meta head and meta neuron atlases. Related to Figure 1. (A) t-SNEs of individual datasets reprocessed and included in our atlases. (B) t-SNE of meta head atlas containing more than 1 million cells/nuclei with cell types defined by unique colours. (C) t-SNEs showing the distribution of cells expressing epithelial (*grh*), muscle (*Mhc*), fat (*AkhR*) and glia (*repo*) marker genes (top), and t-SNEs of the proportional contribution of different source tissue types to each cluster – central brain, optic lobe, and whole head (bottom). (D) t-SNE of meta neuron atlas containing more than 700K cells/nuclei, generated by removing all non-neuronal cells/nuclei from the meta head atlas. (E) t-SNEs showing distribution of cells expressing known marker genes for photoreceptors (*ninaA*), first order interneurons downstream of photoreceptors (*ort*), chemosensory neurons (*Orco*) and mechanosensory neurons (*nompC*) (top), and t-SNEs of the proportional contribution of different source tissue types to each cluster – central brain, optic lobe, and whole head (bottom). (F) t-SNE of meta neuron atlas showing transferred annotation labels from Janssens et al., 2022^36^ to the other datasets (top), and t-SNE of meta neuron atlas showing transferred annotation labels from Ozel et al., 2021^66^ to the other datasets (bottom). Cells without annotation are plotted in white. (G) UMAP (left) and t-SNE (right) of meta central brain neuron atlas containing 329,466 cells/nuclei, generated by removing all non-central brain cells/nuclei from the meta neuron atlas with cell types defined by unique colours.

**Figure S3.**
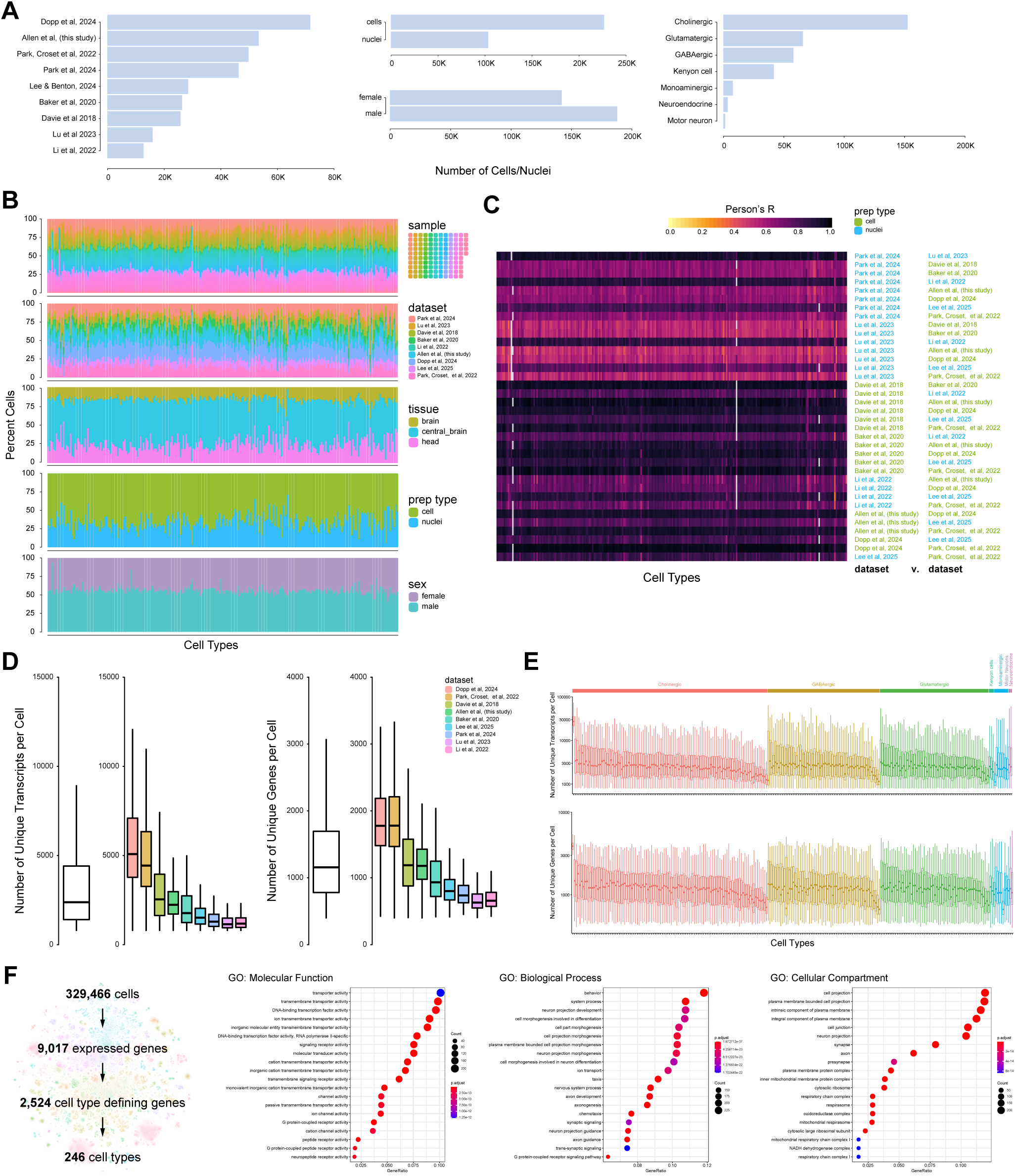
Integration of datasets in central brain neuronal atlas. Related to Figure 1. (A) Bar plots showing the number of cells/nuclei represented for each dataset (left), between cells and nuclei and assigned female or male (middle) and within broad cell type classifications in the meta neuron atlas (right). (B) Stacked bar plots showing the proportional contribution of samples, datasets, tissue, cell preparation, and sex across cell types. (C) Heatmap showing Pearson’s R correlation of dataset versus dataset across all cell types. (D) Boxplot showing the number of unique transcripts per cell (left) across integrated and individual datasets and the number of unique genes detected per cell (right) across integrated and individual datasets. (E) Boxplots displaying the number of unique transcripts (top) and unique genes (bottom) per cell across neuronal cell types. (F) On the left, flow diagram of the central brain neuron atlas, showing the number of single cells profiled, expressed genes (total UMI ≥ 400, max UMI ≥ 4, see STAR Methods), cell type-defining genes (Bonferroni corrected Wilcoxon Signed-Rank Test, log2(FC) > 0.05, padj-value < 0.05), and the number of identified cell types. On the right, gene ontology (GO) enrichment analysis of cell type defining genes for molecular function, biological processes, and cellular compartments.

**Figure S4.**
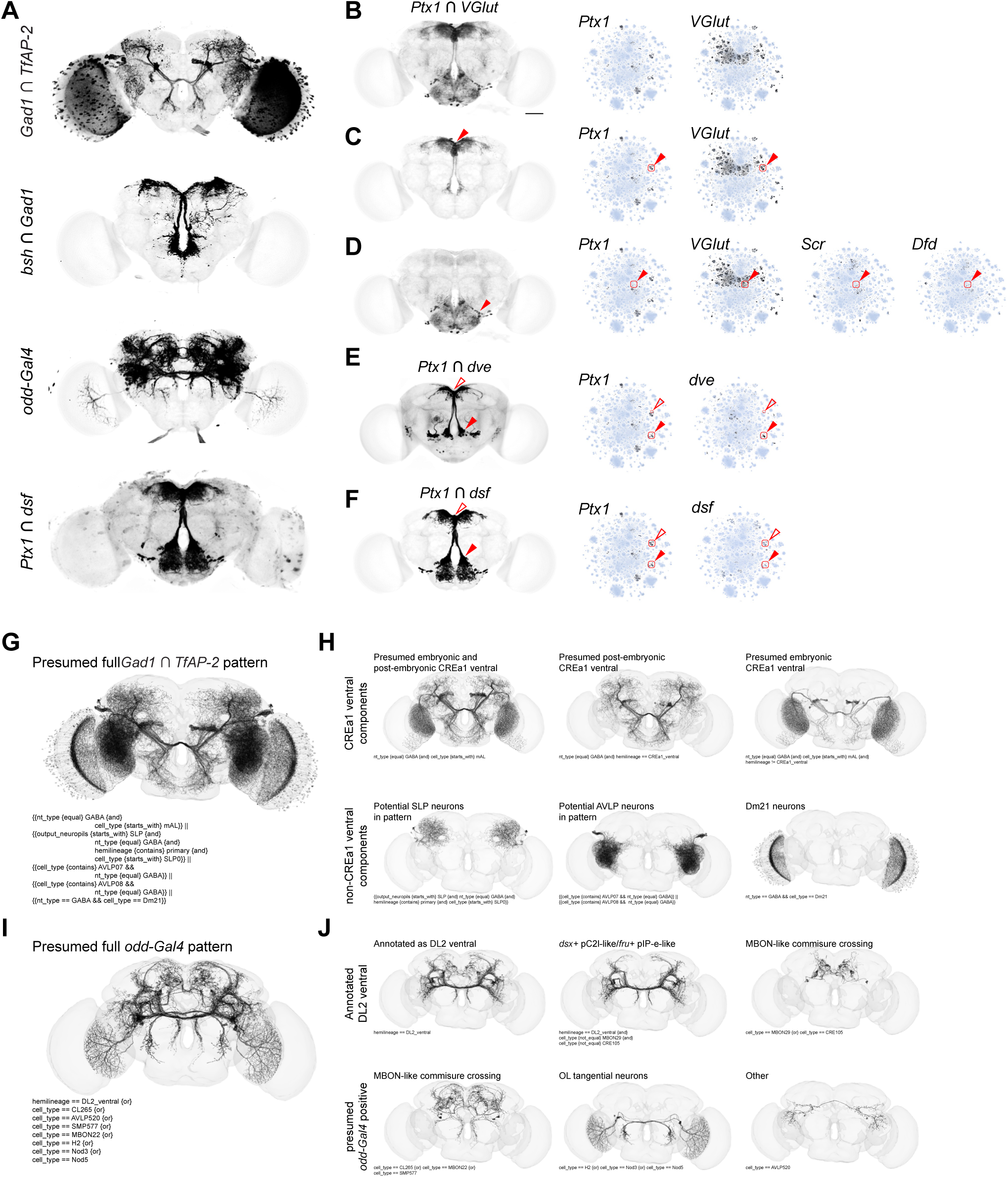
Genetic intersections to identify SMPad1, CREa1 ventral, DM2 central, and DL2 ventral hemilineages. Related to Figure 2. (G) Unsegmented whole-mount immunofluorescence of central brains as seen in Figure 2B. (H) Whole-mount immunofluorescence of central brains showing expression of *VGlut^DBD^*intersected against *Ptx1^AD^* (left). t-SNEs (right) showing *Ptx1* and *VGlut* expression in the central brain neuron atlas. (I) Cerebrum segmented *Ptx1* and *VGlut* intersection expression from panel B (left) and t-SNE plots (right) showing gene *Ptx1* and *VGlut* expression in the central brain neuron atlas. SMPad1 hemilineage cells are highlighted with red box and solid arrowhead. (J) Gnathal ganglia segmented *Ptx1* and *VGlut* intersection expression from panel B (left) and t-SNEs (right) showing gene *Ptx1*, *VGlut*, *Scr*, and *Dfd* expression in the central brain neuron atlas. Gnathal ganglia located cells are highlighted with red box and solid arrowhead. (K) Whole-mount immunofluorescence of central brains showing expression of *dve^DBD^* intersected against *Ptx1^AD^*(left). t-SNEs (right) showing *Ptx1* and *dve* expression in the central brain neuron atlas. Main cell type intersected highlighted with red box and solid arrowhead, subset of SMPad1 hemilineage intersected highlighted with red box and empty red arrow. (L) Whole-mount immunofluorescence of central brains showing expression of *dsf^DBD^* intersected against *Ptx1^AD^* and segmented for clarity (left). t-SNEs (right) showing *Ptx1* and *dsf* expression in the central brain neuron atlas. Main intersected cell types, FLAa3 (red boxes and solid arrowhead) and SMPad1 (red boxes and open arrowhead) hemilineages are highlighted. (M) EM rendering of predicted *Gad1* and *TfAP-2* intersected neurons in the central brain with Flywire identifiers (bellow). (N) EM rendering of predicted CREa1 ventral hemilineage (top). Presumed post-embryonic and embryonic born neurons are shown separately. EM rendering of predicted non-CREa1 ventral components of *Gad1* and *TfAP-2* intersected neurons shown (bottom), including SLP, AVLP, and Dm21 neurons. Flywire identifiers are shown for all cell types. (O) EM rendering of predicted *odd-Gal4* expressing neurons in the central brain with Flywire identifier (bottom). (P) All predicted DL2 ventral neurons with *dsx*+ pC2l-like/*fru*+ pIP-e-like and MBON-like subsets of the hemilineage shown (top). Additional predicted DL2 ventral neurons based on *odd-Gal4* expression, including MBON-like, optic lobe tangential, and other unannotated (bottom). Flywire identifiers are shown for all cell types.

**Figure S5.**
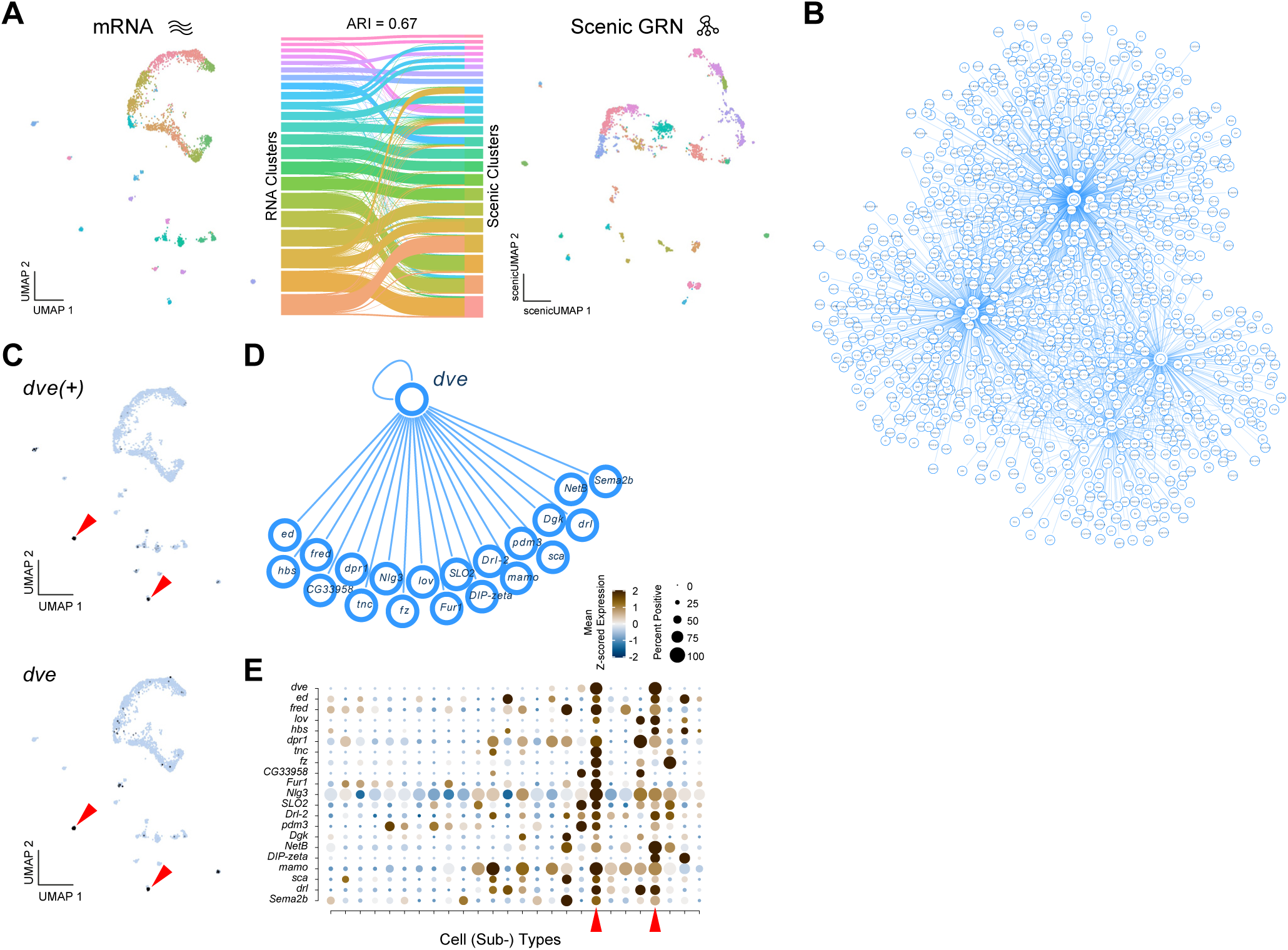
SMPad1 hemilineage gene regulatory network analysis. Related to Figure 2. (A) UMAPs comparison of SMPad1 hemilineage subclustered based on mRNA expression (left) and GRN analysis using SCENIC (right). A Sankey diagram (middle) shows the agreement between mRNA-based and GRN-based clustering, with an adjusted Rand index (ARI) of 0.67. (B) Network representation of SMPad1 hemilineage identified GRNs, where nodes represent genes, and edges indicate regulatory interactions. (C) UMAP showing the expression of the *dve(+)* regulon (top) and *dve* gene expression (bottom) within the SMPad1 hemilineage. (D) Predicted regulatory targets of Dve within the SMPad1 hemilineage based on SCENIC analysis. (E) Dot plot of *dve* and its predicted targets across SMPad1 hemilineage subtypes.

**Figure S6.**
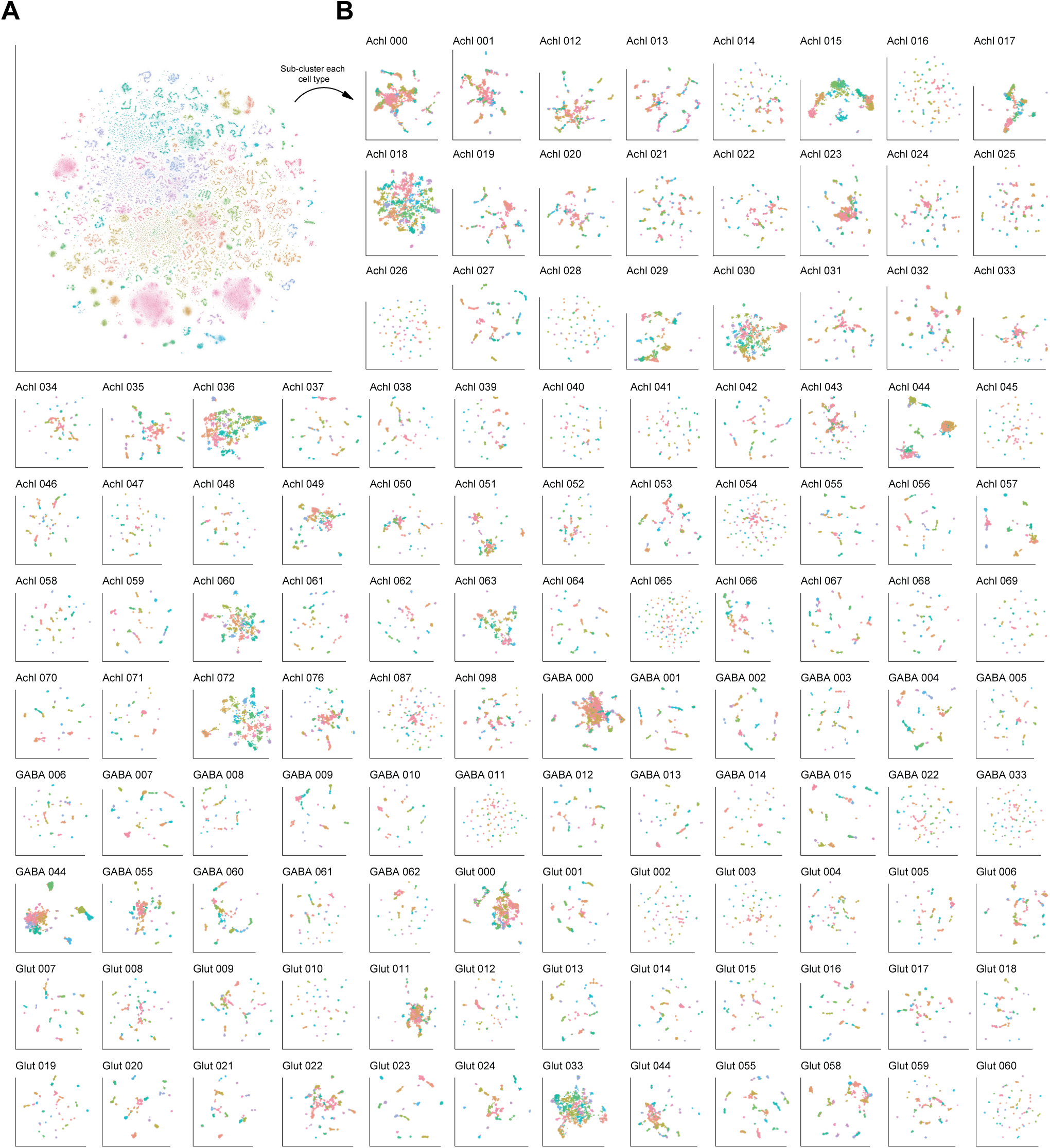
Subclustering of all central brain neuronal cell types. Related to Figure 2. (A) t-SNE of 329,466 central brain neurons representing 9.8x cellular depth coverage, coloured by cell type. (B) Each of the cell types in A was extracted, re-integrated, and re-clustered. UMAPs of the top 120 most numerous cell types (omitting Kenyon cells) are shown, highlighting the thousands of transcriptionally distinct neuronal subtypes that exist in the central brain. The first 20 PCs were used for the UMAP analyses, and Louvain clusters at resolution 2 are shown.

**Figure S7.**
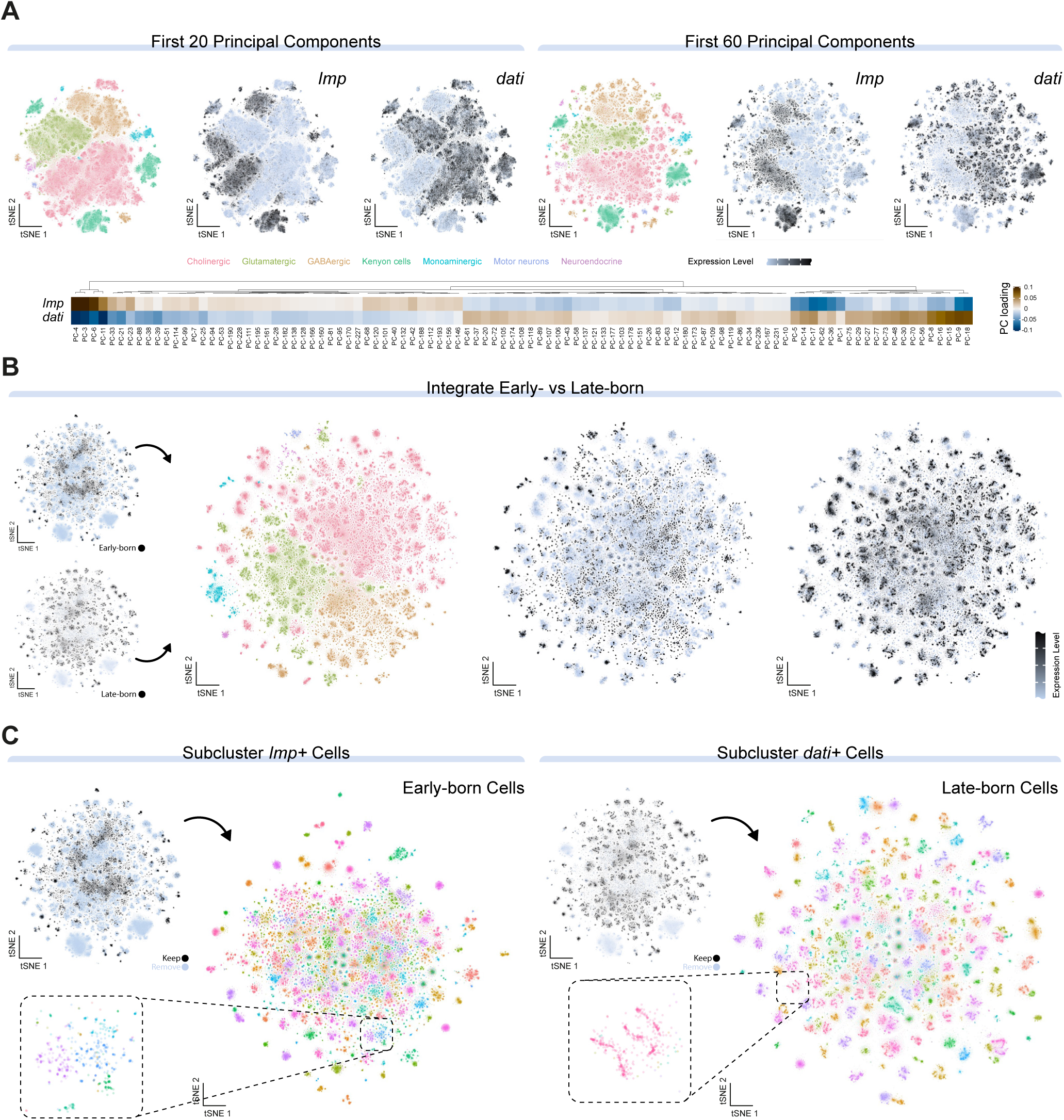
Early- versus late-born neurons exhibit distinct transcriptional profiles. Related to Figure 3. (A) Effects of a varied number of principal components on t-SNE morphology of central brain neurons. (Left) t-SNEs using the first 20 principal components, labelled by broad cell types or *Imp* and *dati* expression. (Right) t-SNEs using the first 60 principal components, labelled by broad cell types or *Imp* and *dati* expression. Separation of the early-born and late-born neurons is constant, and punctate versus serpentine t-SNE morphologies emerge by 60 principal components. Heatmap of top *Imp* and *dati* loadings on principal components (bottom), where at least one had an absolute value greater than 0.01 (bottom), demonstrating *Imp* and *dati’*s contribution to many principal components. (B) Batch correction of early- and late-born neurons. t-SNEs of cells labelled as early-born and late-born before batch correction (left). Integrated clustering of all neuronal cell types following batch correction (middle). Expression of *Imp* and *dati* across the batch corrected atlas (right), showing the transcriptional separation of early- and late-born neurons remains. (C) Sub-clustering of early-born and late-born neuronal populations in the central brain. t-SNEs showing subclusterings of the early-born (*Imp*+) population (left) and late-born (*dati*+) population (right), with insets highlighting representative cluster morphologies. Punctate morphologies in the early-born sub-clustering and serpentine morphologies in the late-born sub-clustering remain.

**Figure S8.**
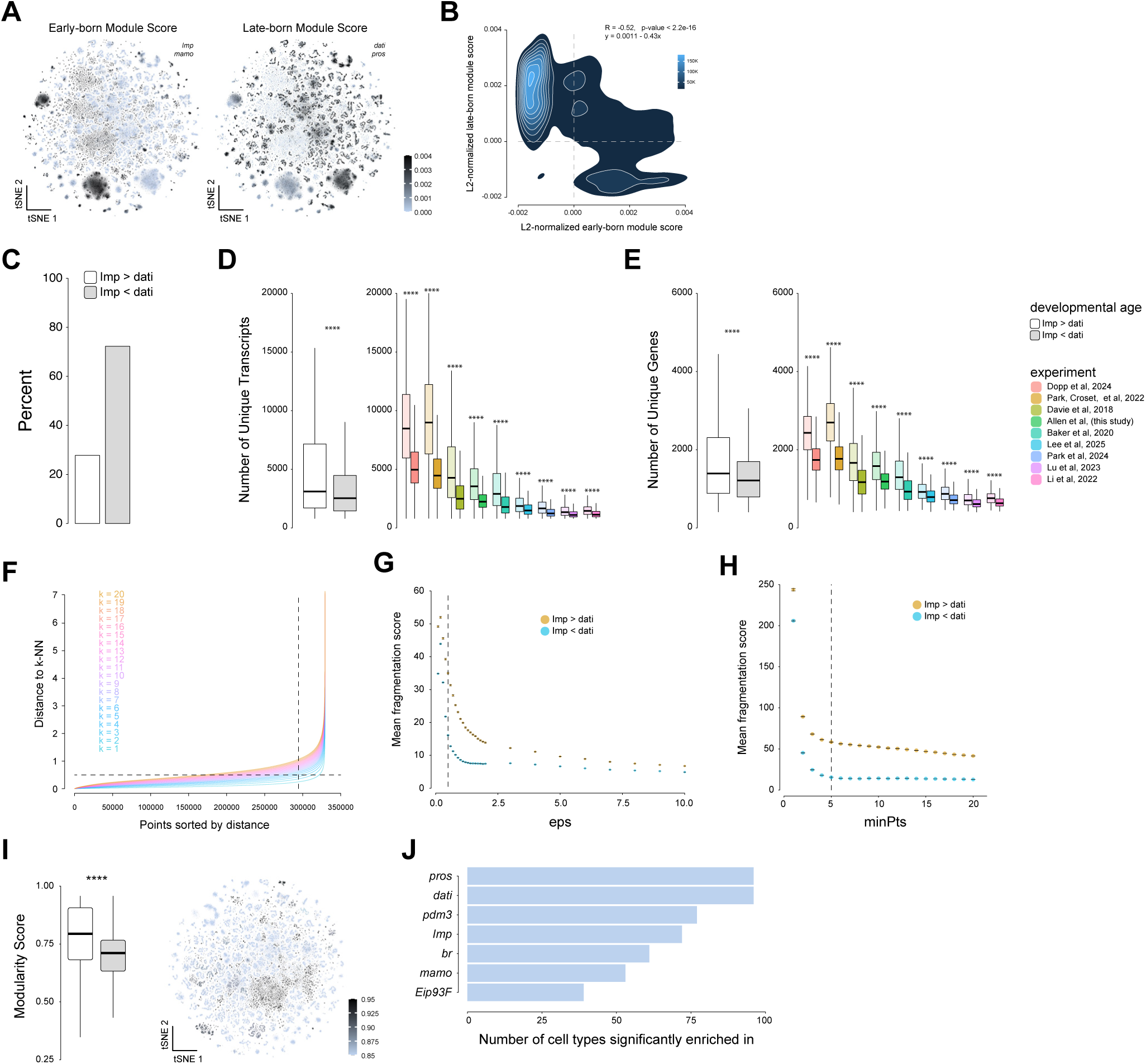
Differing morphologies of early- and late-born neuronal populations. Related to Figure 3. (A) t-SNE of early-born module score (*Imp*+, *mamo+*) and late-born module score (*dati*+, *pros+*) neurons in the adult central brain neuron atlas. (B) A 2D density plot showing the distribution of central brain neurons based on their early-born and late-born module scores. A strong negative correlation is observed between the two scores, indicating that neurons with high expression of early-born module genes tend to show low expression of late-born module genes and vice versa. (C) Bar graph displaying the number of transcriptionally defined cell types significantly enriched for key temporal neurodevelopmental genes. (D) Boxplots showing the number of unique transcripts in early-born and late-born neurons across all (left) or individual (right) datasets. Early-born neurons consistently have more transcripts. (E) Boxplots showing the number of unique genes in early-born and late-born neurons across all (left) or individual (right) datasets. Early-born neurons consistently have more genes. (F) Distance (in t-SNE space) to k^th^-nearest-neighbour (k-NN) for a range of k and sorted distance. Most cells have their 4^th^-NN within a distance of 0.5 (dashed lines). (G) Relationship between the radius of the epsilon neighbourhood (eps) values against the mean fragmentation scores (±95% confidence interval) between early- and late-born cells, showing repeated differences in embedding structure between neuronal types of different developmental age, independent of dbscan parameters. (H) Relationship between the radius of the number of neighbourhood members (minPts) values against the mean fragmentation scores (±95% confidence interval) between early- and late-born cells, showing repeated differences in embedding structure between neuronal types of different developmental age independent of dbscan parameters. (I) Boxplot comparing modularity scores of early- and late-born cells (left) and UMAP visualisation of modularity scores across cell types (right), indicating greater modularity among early-born neurons (Bonferroni corrected Wilcoxon Signed-Rank Test, ****p < 0.0001). (J) Bar graph displaying the number of transcriptionally defined cell types with significant enrichment of key temporal neurodevelopmental genes.

**Figure S9.**
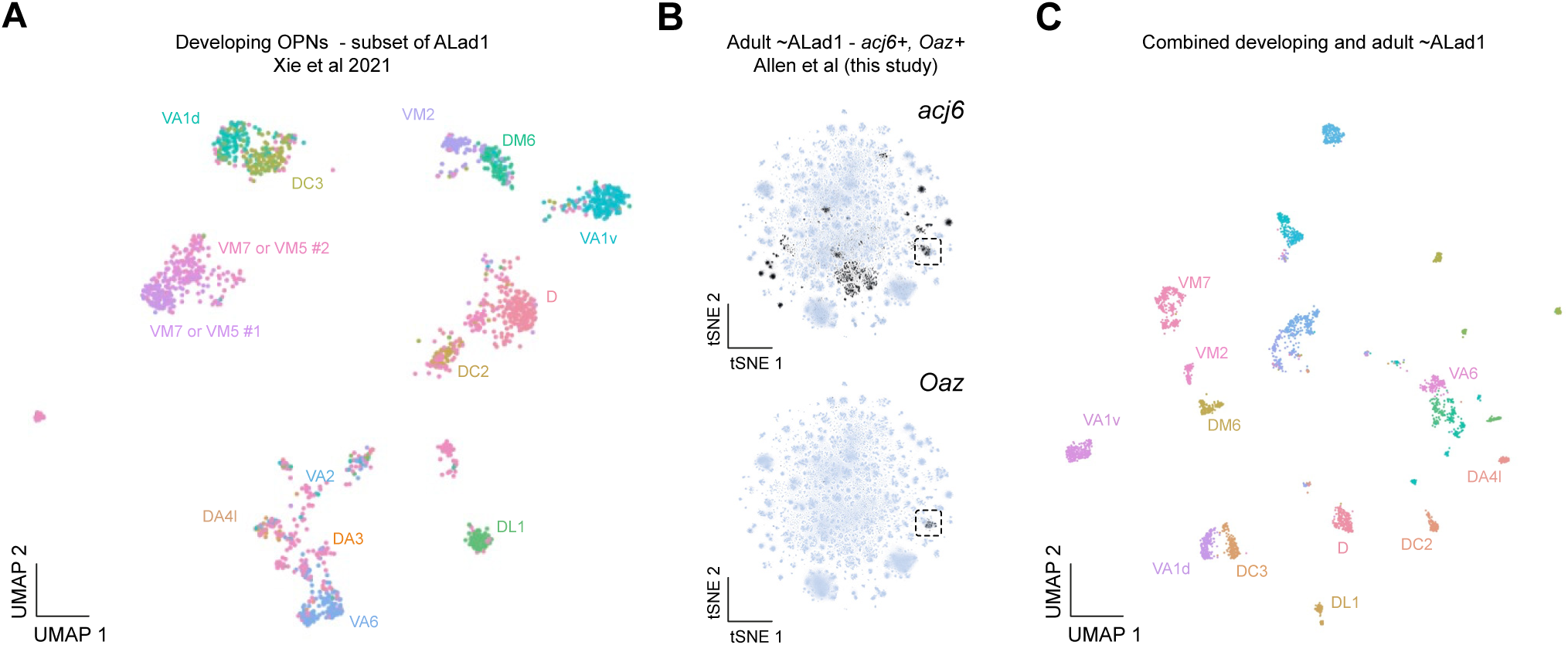
ALad1 hemilineage identification and sub-clustering. Related to Figure 4. (A) UMAP plot of the developing OPN subset of ALad1. Reprocessed data originally from Xie et al., 2021^39^. (B) t-SNE plots showing the ALad1 hemilineage marker genes *acj6* and *Oaz* expression in the central brain neuron atlas. The intersected cells type is highlighted with a dashed black box. (C) UMP plot of combined developing and adult ALad1 hemilineage.

**Figure S10.**
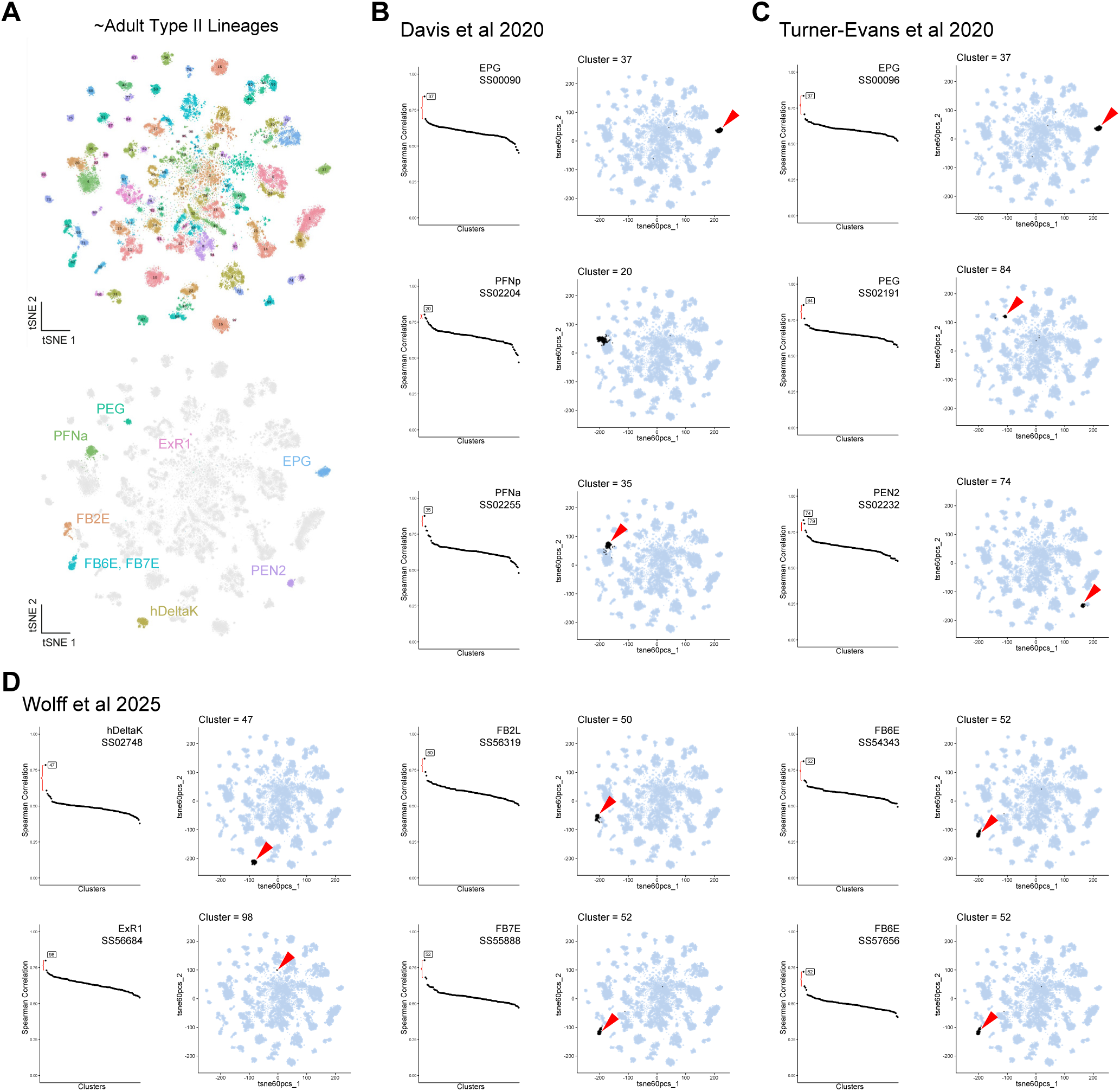
Cell type annotations across the type II neuroblast lineages atlas. Related to Figure 5. (A) t-SNE of type II neuroblast lineage atlas with cell types defined by unique colours and numbers (top) and with cell types annotated based on bulk-seq correlations (bottom). (**B**-**D**). Spearman correlation analysis of cell types in the adult type II atlas compared to available bulk sequencing data, showing the cluster number with the highest correlation (left) and the corresponding cluster in the type II adult atlas (right). Correlations using stable split (SS) labelled, FAC sorted, bulk sequencing data from (**B**) Davis et al., 2020^47^, (**C**) Turner-Evans et al., 2020^48^ and (**D**) Wolff et al., 2025^49^.

**Figure S11.**
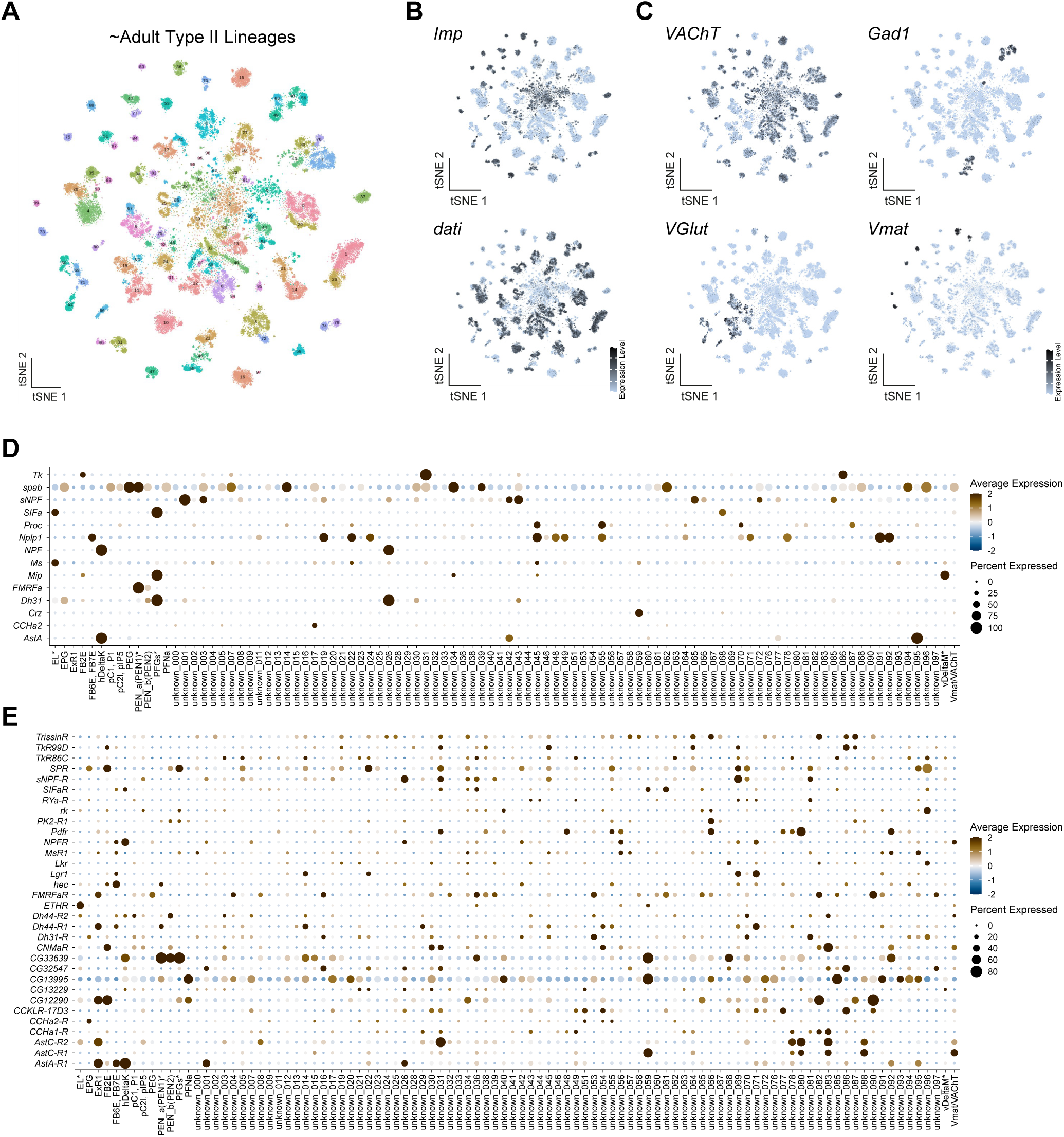
Broad annotations and neuropeptide and neuropeptide receptor expression across the type II neuroblast lineages atlas. Related to Figure 5. (A) t-SNE of re-clustered type II neuroblast lineage atlas with cell types defined by unique colours and numbers. (B) t-SNEs showing the expression of *Imp* (top) and *dati* (bottom) across the type II atlas. (C) t-SNEs showing the expression of neurotransmitter marker genes across the type II atlas. (D) Dot plot of neuropeptide expression across all type II-derived cell types. (E) Dot plot of neuropeptide receptor expression across all type II-derived cell types.

**Figure S12.**
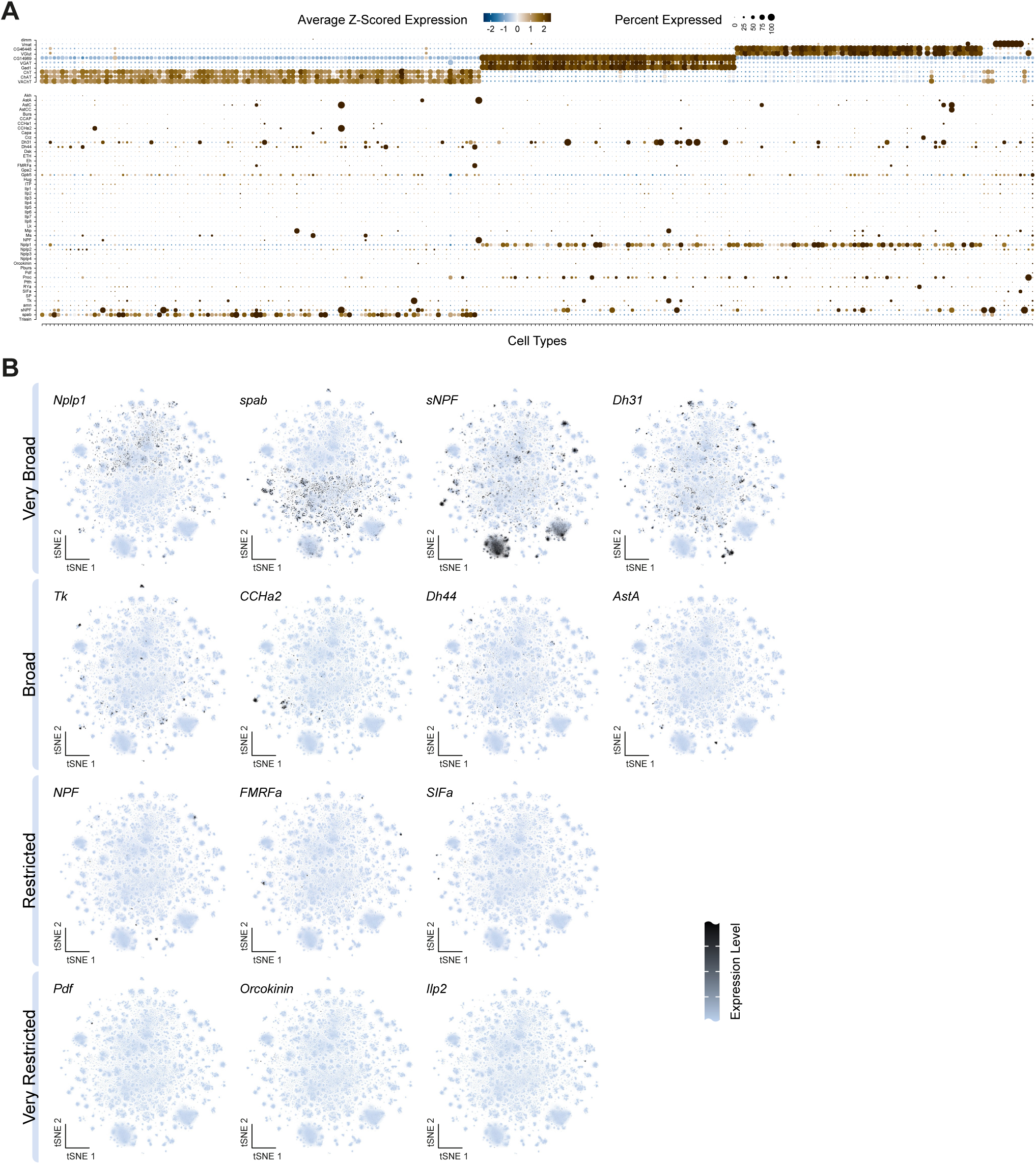
Neuropeptide expression across the central brain. Related to Figure 6. (A) Dot blot of broad cell type identifying genes (above) and all neuropeptide genes (below) across all cell types in the central brain. *Nplp1* is expressed broadly across glutamatergic and GABAergic cell types, while *spab* is expressed broadly across cholinergic cell types. (B) t-SNEs showing examples of categories of neuropeptide gene expression across the central brain: very broadly expressed (*Nplp1*, *spab*, *sNPF*, *Dh31*), broad (*Tk*, *CCHa2*, *Dh44*, *AstA*), restricted (*NPF*, *FMRFa*, *SIFa*) and very restricted (*Pdf*, *Orckokinin*, *Ilp2*).

**Figure S13.**
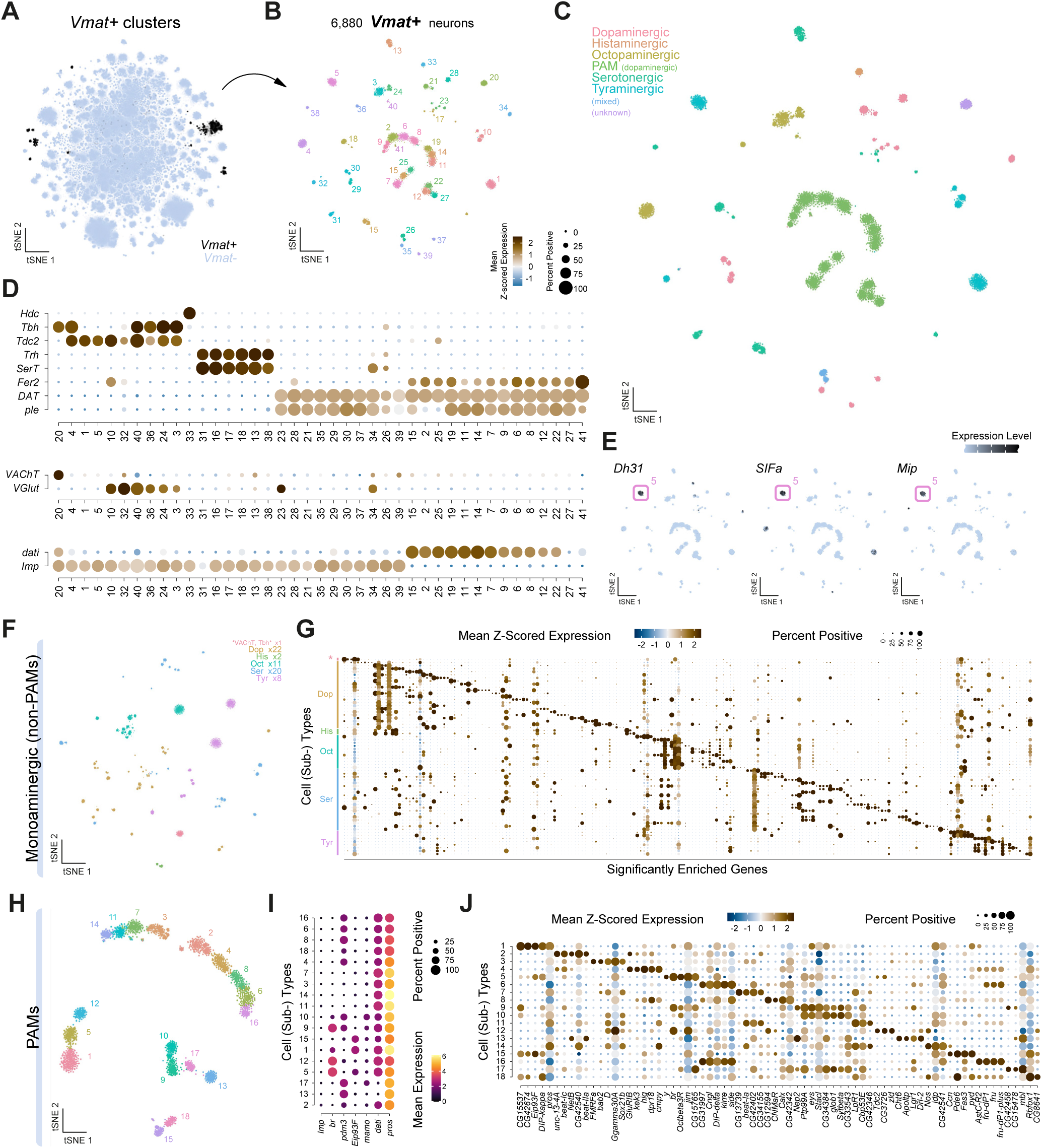
Transcriptional diversity of central brain monoaminergic cell types. Related to Figure 6. (A) t-SNE highlighting monoaminergic cell types based on the expression of the *Vesicular monoamine transporter* (*Vmat*) gene. (B) t-SNE showing sub-clustering analysis of monoaminergic cell types. Transcriptionally distinct cell types are differentially coloured and numbered. (C) Central brain monoaminergic atlas annotated based on monoamine usage. (D) Dot plot of biomarkers across monoaminergic cell types. Monoaminergic markers (top) *Hdc* labels histaminergic neurons, *Tbh* labels octopaminergic neurons, *Tdc-2* labels tyraminergic and octopaminergic neurons, *Trh* and *SerT* label serotonergic neurons, *DAT* and *ple* label dopaminergic neurons. The TF *Fer2* labels the protocerebral anterior medial (PAM) subset of dopamine neurons. Fast-acting neurotransmitter markers (middle) *VAChT* labels cholinergic and *VGlut* glutamatergic neurons. The birth order markers (below) *Imp* labels early-born and *dati* labels late-born neurons. (E) t-SNEs showing example of neuropeptide co-expression of *Dh31*, *SIFa*, and *Mip* in tyraminergic subtype 5. (F) t-SNE showing sub-clustering analysis of monoaminergic cell types with PAMs removed. This analysis identified 64 transcriptionally distinct subtypes. Given the estimated 9.8x depth of coverage of cell types in this atlas, many of these clusters would correspond to subtypes with only 1-3 neurons per hemibrain. (G) Dot plot of top genes defining monoaminergic, non-PAM, cell types. (H) t-SNE showing sub-clustering analysis of dopaminergic PAMs. Transcriptionally distinct cell types are differentially coloured and numbered. This analysis identified 18 transcriptionally distinct subtypes. (I) Dot plot of key developmental birth-order-defining genes across PAM subtypes. (J) Dot plot of top genes defining PAM subtypes.

## Notes

### Competing Interest Statement

The authors have declared no competing interest.

## REFERENCES

1. Davidson, E.H. (2006). The regulatory genome : gene regulatory networks in development and evolution (Academic).

2. Hobert, O., Carrera, I., and Stefanakis, N. (2010). The molecular and gene regulatory signature of a neuron. Trends Neurosci 33, 435–445. 10.1016/j.tins.2010.05.006.

3. Hartenstein, V., and Campos-Ortega, J.A. (1984). Early neurogenesis in wild-type Drosophila melanogaster. Wilehm Roux Arch Dev Biol 193, 308–325. 10.1007/BF00848159.

4. Truman, J.W., and Bate, M. (1988). Spatial and temporal patterns of neurogenesis in the central nervous system of Drosophila melanogaster. Dev Biol 125, 145–157. 10.1016/0012-1606(88)90067-x.

5. Younossi-Hartenstein, A., Nassif, C., Green, P., and Hartenstein, V. (1996). Early neurogenesis of the Drosophila brain. J Comp Neurol 370, 313–329. 10.1002/(SICI)1096-9861(19960701)370:3<313::AID-CNE3>3.0.CO;2-7.

6. Jiang, Y., and Reichert, H. (2014). Drosophila neural stem cells in brain development and tumor formation. J Neurogenet 28, 181–189. 10.3109/01677063.2014.898639.

7. Lee, T. (2017). Wiring the Drosophila Brain with Individually Tailored Neural Lineages. Curr Biol 27, R77–R82. 10.1016/j.cub.2016.12.026.

8. Doe, C.Q. (1992). Molecular markers for identified neuroblasts and ganglion mother cells in the Drosophila central nervous system. Development 116, 855–863. 10.1242/dev.116.4.855.

9. Urbach, R., and Technau, G.M. (2003). Molecular markers for identified neuroblasts in the developing brain of Drosophila. Development 130, 3621–3637. 10.1242/dev.00533.

10. Schlegel, P., Yin, Y., Bates, A.S., Dorkenwald, S., Eichler, K., Brooks, P., Han, D.S., Gkantia, M., Dos Santos, M., Munnelly, E.J., et al. (2024). Whole-brain annotation and multi-connectome cell typing of Drosophila. Nature 634, 139–152. 10.1038/s41586-024-07686-5.

11. Dorkenwald, S., Matsliah, A., Sterling, A.R., Schlegel, P., Yu, S.C., McKellar, C.E., Lin, A., Costa, M., Eichler, K., Yin, Y., et al. (2024). Neuronal wiring diagram of an adult brain. Nature 634, 124–138. 10.1038/s41586-024-07558-y.

12. Holguera, I., and Desplan, C. (2018). Neuronal specification in space and time. Science 362, 176–180. 10.1126/science.aas9435.

13. Pebworth, M.P., Ross, J., Andrews, M., Bhaduri, A., and Kriegstein, A.R. (2021). Human intermediate progenitor diversity during cortical development. Proc Natl Acad Sci U S A 118. 10.1073/pnas.2019415118.

14. Isshiki, T., Pearson, B., Holbrook, S., and Doe, C.Q. (2001). Drosophila Neuroblasts Sequentially Express Transcription Factors which Specify the Temporal Identity of Their Neuronal Progeny. Cell 106, 511–521. 10.1016/s0092-8674(01)00465-2.

15. Bayraktar, O.A., and Doe, C.Q. (2013). Combinatorial temporal patterning in progenitors expands neural diversity. Nature 498, 449–455. 10.1038/nature12266.

16. Barabasi, D.L., Ferreira Castro, A., and Engert, F. (2025). Three systems of circuit formation: assembly, updating and tuning. Nat Rev Neurosci 26, 232–243. 10.1038/s41583-025-00910-9.

17. Croset, V., Treiber, C.D., and Waddell, S. (2018). Cellular diversity in the Drosophila midbrain revealed by single-cell transcriptomics. Elife 7. 10.7554/eLife.34550.

18. Park, A., Croset, V., Otto, N., Agarwal, D., Treiber, C.D., Meschi, E., Sims, D., and Waddell, S. (2022). Gliotransmission of D-serine promotes thirst-directed behaviors in Drosophila. Curr Biol 32, 3952–3970 e3958. 10.1016/j.cub.2022.07.038.

19. Dopp, J., Ortega, A., Davie, K., Poovathingal, S., Baz, E.S., and Liu, S. (2024). Single-cell transcriptomics reveals that glial cells integrate homeostatic and circadian processes to drive sleep-wake cycles. Nat Neurosci 27, 359–372. 10.1038/s41593-023-01549-4.

20. Allen, A.M., Neville, M.C., Nojima, T., Alejevski, F., and Goodwin, S.F. (2025). A Role for Exaptation in Sculpting Sexually Dimorphic Brains from Shared Neural Lineages.

21. Li, H., Janssens, J., De Waegeneer, M., Kolluru, S.S., Davie, K., Gardeux, V., Saelens, W., David, F.P.A., Brbic, M., Spanier, K., et al. (2022). Fly Cell Atlas: A single-nucleus transcriptomic atlas of the adult fruit fly. Science 375, eabk2432. 10.1126/science.abk2432.

22. Yeung, K., Bollepogu Raja, K.K., Shim, Y.K., Li, Y., Chen, R., and Mardon, G. (2022). Single cell RNA sequencing of the adult Drosophila eye reveals distinct clusters and novel marker genes for all major cell types. Commun Biol 5, 1370. 10.1038/s42003-022-04337- 1.

23. Ito, K., Shinomiya, K., Ito, M., Armstrong, J.D., Boyan, G., Hartenstein, V., Harzsch, S., Heisenberg, M., Homberg, U., Jenett, A., et al. (2014). A systematic nomenclature for the insect brain. Neuron 81, 755–765. 10.1016/j.neuron.2013.12.017.

24. Allen, A.M., Neville, M.C., Birtles, S., Croset, V., Treiber, C.D., Waddell, S., and Goodwin, S.F. (2020). A single-cell transcriptomic atlas of the adult Drosophila ventral nerve cord. Elife 9. 10.7554/eLife.54074.

25. Diao, F., Vasudevan, D., Heckscher, E.S., and White, B.H. (2024). Hox gene-specific cellular targeting using split intein Trojan exons. Proc Natl Acad Sci U S A 121, e2317083121. 10.1073/pnas.2317083121.

26. Hirth, F., Hartmann, B., and Reichert, H. (1998). Homeotic gene action in embryonic brain development of Drosophila. Development 125, 1579–1589. 10.1242/dev.125.9.1579.

27. Holland, P.W. (2013). Evolution of homeobox genes. Wiley Interdiscip Rev Dev Biol 2, 31–45. 10.1002/wdev.78.

28. Marin, E.C., Morris, B.J., Stürner, T., Champion, A.S., Krzeminski, D., Badalamente, G., Gkantia, M., Dunne, C.R., Eichler, K., Takemura, S.-Y., et al. (2024). Systematic annotation of a complete adult male Drosophila nerve cord connectome reveals principles of functional organisation. Elife 13.

29. McGinnis, O.J., Alejevski, F., Neville, M.C., Goodwin, S.F., and Allen, A.M. (2025). Transcriptomic identity and diveristy of cell-types of the Drosophila navigation complex. (in prep).

30. Soffers, J.H., Beck, E., Sytkowski, D.J., Maughan, M.E., Devasri, D., Zhu, Y., Wilson, B., Chen, Y.D., Erclik, T., Truman, J.W., et al. (2025). A library of lineage-specific driver lines connects developing neuronal circuits to behavior in the Drosophila Ventral Nerve Cord. Elife 14. 10.7554/eLife.106042.1.

31. Aibar, S., Gonzalez-Blas, C.B., Moerman, T., Huynh-Thu, V.A., Imrichova, H., Hulselmans, G., Rambow, F., Marine, J.C., Geurts, P., Aerts, J., et al. (2017). SCENIC: single-cell regulatory network inference and clustering. Nat Methods 14, 1083–1086. 10.1038/nmeth.4463.

32. Van de Sande, B., Flerin, C., Davie, K., De Waegeneer, M., Hulselmans, G., Aibar, S., Seurinck, R., Saelens, W., Cannoodt, R., Rouchon, Q., et al. (2020). A scalable SCENIC workflow for single-cell gene regulatory network analysis. Nat Protoc 15, 2247–2276. 10.1038/s41596-020-0336-2.

33. Maurange, C., Cheng, L., and Gould, A.P. (2008). Temporal transcription factors and their targets schedule the end of neural proliferation in Drosophila. Cell 133, 891–902. 10.1016/j.cell.2008.03.034.

34. Yang, C.P., Samuels, T.J., Huang, Y., Yang, L., Ish-Horowicz, D., Davis, I., and Lee, T. (2017). Imp and Syp RNA-binding proteins govern decommissioning of Drosophila neural stem cells. Development 144, 3454–3464. 10.1242/dev.149500.

35. Etheredge, J. (2017). Transcriptional profiling of Drosophila larval ventral nervous system hemilineages. Doctoral Thesis (University of Cambridge).

36. Janssens, J., Aibar, S., Taskiran, II, Ismail, J.N., Gomez, A.E., Aughey, G., Spanier, K.I., De Rop, F.V., Gonzalez-Blas, C.B., Dionne, M., et al. (2022). Decoding gene regulation in the fly brain. Nature 601, 630–636. 10.1038/s41586-021-04262-z.

37. Zhou, B., Williams, D.W., Altman, J., Riddiford, L.M., and Truman, J.W. (2009). Temporal patterns of broad isoform expression during the development of neuronal lineages in Drosophila. Neural Dev 4, 39. 10.1186/1749-8104-4-39.

38. Yu, H.H., Kao, C.F., He, Y., Ding, P., Kao, J.C., and Lee, T. (2010). A complete developmental sequence of a Drosophila neuronal lineage as revealed by twin-spot MARCM. PLoS Biol 8. 10.1371/journal.pbio.1000461.

39. Xie, Q., Brbic, M., Horns, F., Kolluru, S.S., Jones, R.C., Li, J., Reddy, A.R., Xie, A., Kohani, S., Li, Z., et al. (2021). Temporal evolution of single-cell transcriptomes of Drosophila olfactory projection neurons. Elife 10. 10.7554/eLife.63450.

40. Kohwi, M., Hiebert, L.S., and Doe, C.Q. (2011). The pipsqueak-domain proteins Distal antenna and Distal antenna-related restrict Hunchback neuroblast expression and early- born neuronal identity. Development 138, 1727–1735. 10.1242/dev.061499.

41. Li, Q., Barish, S., Okuwa, S., Maciejewski, A., Brandt, A.T., Reinhold, D., Jones, C.D., and Volkan, P.C. (2016). A Functionally Conserved Gene Regulatory Network Module Governing Olfactory Neuron Diversity. PLoS Genet 12, e1005780. 10.1371/journal.pgen.1005780.

42. Sousa-Nunes, R., Cheng, L.Y., and Gould, A.P. (2010). Regulating neural proliferation in the Drosophila CNS. Curr Opin Neurobiol 20, 50–57. 10.1016/j.conb.2009.12.005.

43. Homem, C.C., Repic, M., and Knoblich, J.A. (2015). Proliferation control in neural stem and progenitor cells. Nat Rev Neurosci 16, 647–659. 10.1038/nrn4021.

44. Doe, C.Q. (2017). Temporal Patterning in the Drosophila CNS. Annu Rev Cell Dev Biol 33, 219–240. 10.1146/annurev-cellbio-111315-125210.

45. Michki, N.S., Li, Y., Sanjasaz, K., Zhao, Y., Shen, F.Y., Walker, L.A., Cao, W., Lee, C.Y., and Cai, D. (2021). The molecular landscape of neural differentiation in the developing Drosophila brain revealed by targeted scRNA-seq and multi-informatic analysis. Cell Rep 35, 109039.v 10.1016/j.celrep.2021.109039.

46. Rajan, A., Anhezini, L., Rives-Quinto, N., Chhabra, J.Y., Neville, M.C., Larson, E.D., Goodwin, S.F., Harrison, M.M., and Lee, C.Y. (2023). Low-level repressive histone marks fine-tune gene transcription in neural stem cells. Elife 12. 10.7554/eLife.86127.

47. Davis, F.P., Nern, A., Picard, S., Reiser, M.B., Rubin, G.M., Eddy, S.R., and Henry, G.L. (2020). A genetic, genomic, and computational resource for exploring neural circuit function. Elife 9. 10.7554/eLife.50901.

48. Turner-Evans, D.B., Jensen, K.T., Ali, S., Paterson, T., Sheridan, A., Ray, R.P., Wolff, T., Lauritzen, J.S., Rubin, G.M., Bock, D.D., and Jayaraman, V. (2020). The Neuroanatomical Ultrastructure and Function of a Biological Ring Attractor. Neuron 108, 145–163 e110. 10.1016/j.neuron.2020.08.006.

49. Wolff, T., Eddison, M., Chen, N., Nern, A., Sundaramurthi, P., Sitaraman, D., and Rubin, G.M. (2025). Cell type-specific driver lines targeting the Drosophila central complex and their use to investigate neuropeptide expression and sleep regulation. ELife 14.

50. Hulse, B.K., Haberkern, H., Franconville, R., Turner-Evans, D., Takemura, S.-Y., Wolff, T., Noorman, M., Dreher, M., Dan, C., Parekh, R., et al. (2021). A connectome of the *Drosophila* central complex reveals network motifs suitable for flexible navigation and context-dependent action selection eLife *10*, e66039 , citation = eLife 62021;66010 e66039. 10.7554/eLife.66039.

51. Heinze, S. (2023). The Insect Central Complex. Oxford University Press.

52. Nassel, D.R. (2025). What Drosophila can tell us about state-dependent peptidergic signaling in insects. Insect Biochem Mol Biol 179, 104275. 10.1016/j.ibmb.2025.104275.

53. Hamanaka, Y., Park, D., Yin, P., Annangudi, S.P., Edwards, T.N., Sweedler, J., Meinertzhagen, I.A., and Taghert, P.H. (2010). Transcriptional orchestration of the regulated secretory pathway in neurons by the bHLH protein DIMM. Curr Biol 20, 9–18. 10.1016/j.cub.2009.11.065.

54. Hewes, R.S., Park, D., Gauthier, S.A., Schaefer, A.M., and Taghert, P.H. (2003). The bHLH protein Dimmed controls neuroendocrine cell differentiation in Drosophila. Development 130, 1771–1781. 10.1242/dev.00404.

55. Park, D., Veenstra, J.A., Park, J.H., and Taghert, P.H. (2008). Mapping peptidergic cells in Drosophila: where DIMM fits in. PLoS One 3, e1896. 10.1371/journal.pone.0001896.

56. Nassel, D.R., and Zandawala, M. (2022). Endocrine cybernetics: neuropeptides as molecular switches in behavioural decisions. Open Biol 12, 220174. 10.1098/rsob.220174.

57. Melcher, C., and Pankratz, M.J. (2005). Candidate gustatory interneurons modulating feeding behavior in the Drosophila brain. PLoS Biol 3, e305. 10.1371/journal.pbio.0030305.

58. Bader, R., Colomb, J., Pankratz, B., Schrock, A., Stocker, R.F., and Pankratz, M.J. (2007). Genetic dissection of neural circuit anatomy underlying feeding behavior in Drosophila: distinct classes of hugin-expressing neurons. J Comp Neurol 502, 848–856. 10.1002/cne.21342.

59. King, A.N., Barber, A.F., Smith, A.E., Dreyer, A.P., Sitaraman, D., Nitabach, M.N., Cavanaugh, D.J., and Sehgal, A. (2017). A Peptidergic Circuit Links the Circadian Clock to Locomotor Activity. Curr Biol 27, 1915–1927 e1915. 10.1016/j.cub.2017.05.089.

60. Sterne, G.R., Otsuna, H., Dickson, B.J., and Scott, K. (2021). Classification and genetic targeting of cell types in the primary taste and premotor center of the adult Drosophila brain. Elife 10. 10.7554/eLife.71679.

61. Schwarz, J.E., King, A.N., Hsu, C.T., Barber, A.F., and Sehgal, A. (2021). Hugin(+) neurons provide a link between sleep homeostat and circadian clock neurons. Proc Natl Acad Sci U S A 118. 10.1073/pnas.2111183118.

62. Holder, B.L., and Dissel, S. (2025). Cell-specific tools for understanding behavior. Elife 14. 10.7554/eLife.106686.

63. Larsen, C., Shy, D., Spindler, S.R., Fung, S., Pereanu, W., Younossi-Hartenstein, A., and Hartenstein, V. (2009). Patterns of growth, axonal extension and axonal arborization of neuronal lineages in the developing Drosophila brain. Dev Biol 335, 289–304. 10.1016/j.ydbio.2009.06.015.

64. Lee, Y.J., Yang, C.P., Miyares, R.L., Huang, Y.F., He, Y., Ren, Q., Chen, H.M., Kawase, T., Ito, M., Otsuna, H., et al. (2020). Conservation and divergence of related neuronal lineages in the Drosophila central brain. Elife 9. 10.7554/eLife.53518.

65. Walsh, K.T., and Doe, C.Q. (2017). Drosophila embryonic type II neuroblasts: origin, temporal patterning, and contribution to the adult central complex. Development 144, 4552–4562. 10.1242/dev.157826.

66. Ozel, M.N., Simon, F., Jafari, S., Holguera, I., Chen, Y.C., Benhra, N., El-Danaf, R.N., Kapuralin, K., Malin, J.A., Konstantinides, N., and Desplan, C. (2021). Neuronal diversity and convergence in a visual system developmental atlas. Nature 589, 88–95. 10.1038/s41586-020-2879-3.

67. Davie, K., Janssens, J., Koldere, D., De Waegeneer, M., Pech, U., Kreft, L., Aibar, S., Makhzami, S., Christiaens, V., Bravo Gonzalez-Blas, C., et al. (2018). A Single-Cell Transcriptome Atlas of the Aging Drosophila Brain. Cell 174, 982–998 e920. 10.1016/j.cell.2018.05.057.

68. Baker, B.M., Mokashi, S.S., Shankar, V., Hatfield, J.S., Hannah, R.C., Mackay, T.F.C., and Anholt, R.R.H. (2021). The Drosophila brain on cocaine at single-cell resolution. Genome Res 31, 1927–1937. 10.1101/gr.268037.120.

69. Lee, D., Shahandeh, M.P., Abuin, L., and Benton, R. (2025). Comparative single-cell transcriptomic atlases of drosophilid brains suggest glial evolution during ecological adaptation. PLoS Biol 23, e3003120. 10.1371/journal.pbio.3003120.

70. Kurmangaliyev, Y.Z., Yoo, J., Valdes-Aleman, J., Sanfilippo, P., and Zipursky, S.L. (2020). Transcriptional Programs of Circuit Assembly in the Drosophila Visual System. Neuron 108, 1045–1057 e1046. 10.1016/j.neuron.2020.10.006.

71. Lu, T.C., Brbic, M., Park, Y.J., Jackson, T., Chen, J., Kolluru, S.S., Qi, Y., Katheder, N.S., Cai, X.T., Lee, S., et al. (2023). Aging Fly Cell Atlas identifies exhaustive aging features at cellular resolution. Science 380, eadg0934. 10.1126/science.adg0934.

72. Park, Y.J., Lu, T.C., Jackson, T., Goodman, L.D., Ran, L., Chen, J., Liang, C.Y., Harrison, E., Ko, C., Hsu, A.L., et al. (2024). Whole organism snRNA-seq reveals systemic peripheral changes in Alzheimer’s Disease fly models. bioRxiv. 10.1101/2024.03.10.584317.

73. Konstantinides, N., Kapuralin, K., Fadil, C., Barboza, L., Satija, R., and Desplan, C. (2018). Phenotypic Convergence: Distinct Transcription Factors Regulate Common Terminal Features. Cell 174, 622–635 e613. 10.1016/j.cell.2018.05.021.

74. Rideout, E.J., Dornan, A.J., Neville, M.C., Eadie, S., and Goodwin, S.F. (2010). Control of sexual differentiation and behavior by the doublesex gene in Drosophila melanogaster. Nat Neurosci 13, 458–466. 10.1038/nn.2515.

75. Soneson, C., and Srivastava, A. (2021). alevinQC: Generate QC Reports For Alevin Output.

76. Pagès, H., Carlson, M., Falcon, S., and Li, N. (2020). AnnotationDbi: Manipulation of SQLite-based annotations in Bioconductor.

77. Sims, D., Ilott, N.E., Sansom, S.N., Sudbery, I.M., Johnson, J.S., Fawcett, K.A., Berlanga- Taylor, A.J., Luna-Valero, S., Ponting, C.P., and Heger, A. (2014). CGAT: computational genomics analysis toolkit. Bioinformatics 30, 1290–1291. 10.1093/bioinformatics/btt756.

78. Yu, G., Wang, L.G., Han, Y., and He, Q.Y. (2012). clusterProfiler: an R package for comparing biological themes among gene clusters. OMICS 16, 284–287. 10.1089/omi.2011.0118.

79. Gu, Z., Eils, R., and Schlesner, M. (2016). Complex heatmaps reveal patterns and correlations in multidimensional genomic data. Bioinformatics 32, 2847–2849. 10.1093/bioinformatics/btw313.

80. Kuhn, M., Jackson, S., and Cimentada, J. (2020). corrr: Correlations in R.

81. Wilke, C.O. (2020). cowplot: Streamlined Plot Theme and Plot Annotations for ’ggplot2’.

82. Shannon, P., Markiel, A., Ozier, O., Baliga, N.S., Wang, J.T., Ramage, D., Amin, N., Schwikowski, B., and Ideker, T. (2003). Cytoscape: a software environment for integrated models of biomolecular interaction networks. Genome Res 13, 2498–2504. 10.1101/gr.1239303.

83. McGinnis, C.S., Murrow, L.M., and Gartner, Z.J. (2019). DoubletFinder: Doublet Detection in Single-Cell RNA Sequencing Data Using Artificial Nearest Neighbors. Cell Syst 8, 329–337 e324. 10.1016/j.cels.2019.03.003.

84. Wickham, H., François, R., Henry, L., Müller, K., and Vaughan, D. (2021). dplyr: A Grammar of Data Manipulation.

85. Lun, A.T.L., Riesenfeld, S., Andrews, T., Dao, T.P., Gomes, T., participants in the 1st 1532 Human Cell Atlas, J., and Marioni, J.C. (2019). EmptyDrops: distinguishing cells from empty droplets in droplet-based single-cell RNA sequencing data. Genome Biol 20, 63. 10.1186/s13059-019-1662-y.

86. Jenkins, V.K., Larkin, A., Thurmond, J., and FlyBase, C. (2022). Using FlyBase: A Database of Drosophila Genes and Genetics. Methods Mol Biol 2540, 1-34. 10.1007/978-1-0716-2541-5_1.

84. Bengtsson, H. (2021). A unifying framework for parallel and distributed processing in R using futures. The R Journal 13, 208–291.

85. Kassambara, A. (2019). ggcorrplot: Visualization of a Correlation Matrix using ’ggplot2’.

86. Wickham, H. (2016). ggplot2: Elegant Graphics for Data Analysis.

87. Kassambara, A. (2020). ggpubr: ’ggplot2’ Based Publication Ready Plots.

88. Korsunsky, I., Millard, N., Fan, J., Slowikowski, K., Zhang, F., Wei, K., Baglaenko, Y., Brenner, M., Loh, P.R., and Raychaudhuri, S. (2019). Fast, sensitive and accurate integration of single-cell data with Harmony. Nat Methods 16, 1289–1296. 10.1038/s41592-019-0619-0.

89. Cao, J., Spielmann, M., Qiu, X., Huang, X., Ibrahim, D.M., Hill, A.J., Zhang, F., Mundlos, S., Christiansen, L., Steemers, F.J., et al. (2019). The single-cell transcriptional landscape of mammalian organogenesis. Nature 566, 496–502. 10.1038/s41586-019-0969-x.

90. Carlson, M. (2020). org.Dm.eg.db: Genome wide annotation for Fly.

91. Pedersen, T. L. (2020). patchwork: The Composer of Plots.

92. Kolde, R. (2019). pheatmap: Pretty Heatmaps.

93. Wickham, H., and Hester, J. (2020). readr: Read Rectangular Text Data.

94. Goodstadt, L. (2010). Ruffus: a lightweight Python library for computational pipelines. Bioinformatics 26, 2778–2779. 10.1093/bioinformatics/btq524.

95. Wolock, S.L., Lopez, R., and Klein, A.M. (2019). Scrublet: Computational Identification of Cell Doublets in Single-Cell Transcriptomic Data. Cell Syst 8, 281–291 e289. 10.1016/j.cels.2018.11.005.

96. Hao, Y., Hao, S., Andersen-Nissen, E., Mauck, W.M., 3rd, Zheng, S., Butler, A., Lee, M.J., Wilk, A.J., Darby, C., Zager, M., et al. (2021). Integrated analysis of multimodal single-cell data. Cell *184*, 3573-3587 e3529. 10.1016/j.cell.2021.04.048.

97. Satija, R., Butler, A., Hoffman, P., and Stuart, T. (2021). SeuratObject: Data Structures for Single Cell Data.

98. Chang, W., Cheng, J., Allaire, J.J., Sievert, C., Schloerke, B., Xie, Y., Allen, J., McPherson, J., Dipert, A., and Borges, B. (2022). shiny: Web Application Framework for R.

99. Ouyang, J.F., Kamaraj, U.S., Cao, E.Y., and Rackham, O.J.L. (2021). ShinyCell: simple and sharable visualization of single-cell gene expression data. Bioinformatics 37, 3374–3376. 10.1093/bioinformatics/btab209.

100. Amezquita, R.A., Lun, A.T.L., Becht, E., Carey, V.J., Carpp, L.N., Geistlinger, L., Marini, F., Rue-Albrecht, K., Risso, D., Soneson, C., et al. (2020). Orchestrating single-cell analysis with Bioconductor. Nat Methods 17, 137–145. 10.1038/s41592-019-0654-x.

101. Bernstein, N.J., Fong, N.L., Lam, I., Roy, M.A., Hendrickson, D.G., and Kelley, D.R. (2020). Solo: Doublet Identification in Single-Cell RNA-Seq via Semi-Supervised Deep Learning. Cell Syst 11, 95–101 e105. 10.1016/j.cels.2020.05.010.

102. Young, M.D., and Behjati, S. (2020). SoupX removes ambient RNA contamination from droplet-based single-cell RNA sequencing data. Gigascience 9. 10.1093/gigascience/giaa151.

103. Wickham, H. (2019). stringr: Simple, Consistent Wrappers for Common String Operations.

104. Morgan, M., Obenchain, V., Hester, J., and Pagès, H. (2020). SummarizedExperiment: SummarizedExperiment container.

105. Müller, K., and Wickham, H. (2021). tibble: Simple Data Frames.

106. Wickham, H. (2021). tidyr: Tidy Messy Data.

107. Wan, Y., Otsuna, H., Holman, H.A., Bagley, B., Ito, M., Lewis, A.K., Colasanto, M., Kardon, G., Ito, K., and Hansen, C. (2017). FluoRender: joint freehand segmentation and visualization for many-channel fluorescence data analysis. BMC Bioinformatics 18. 10.1186/s12859-017-1694-9.

108. Lillvis, J.L., Otsuna, H., Ding, X., Pisarev, I., Kawase, T., Colonell, J., Rokicki, K., Goina, C., Gao, R., Hu, A., et al. (2022). Rapid reconstruction of neural circuits using tissue expansion and light sheet microscopy. eLife 11. 10.7554/elife.81248.

109. Zeileis, A., and Grothendieck, G. (2005). zoo: S3 Infrastructure for Regular and Irregular Time Series. Journal of Statistical Software 14, 1–27. 10.18637/jss.v014.i06.

110. Srivastava, A., Malik, L., Smith, T., Sudbery, I., and Patro, R. (2019). Alevin efficiently estimates accurate gene abundances from dscRNA-seq data. Genome Biol 20, 65. 10.1186/s13059-019-1670-y.

111. Nagarkar-Jaiswal, S., DeLuca, S.Z., Lee, P.T., Lin, W.W., Pan, H., Zuo, Z., Lv, J., Spradling, A.C., and Bellen, H.J. (2015). A genetic toolkit for tagging intronic MiMIC containing genes. Elife 4. 10.7554/eLife.08469.

112. Lee, P.T., Zirin, J., Kanca, O., Lin, W.W., Schulze, K.L., Li-Kroeger, D., Tao, R., Devereaux, C., Hu, Y., Chung, V., et al. (2018). A gene-specific T2A-GAL4 library for Drosophila. Elife 7. 10.7554/eLife.35574.

113. Diao, F., Ironfield, H., Luan, H., Diao, F., Shropshire, W.C., Ewer, J., Marr, E., Potter, C.J., Landgraf, M., and White, B.H. (2015). Plug-and-play genetic access to drosophila cell types using exchangeable exon cassettes. Cell Rep 10, 1410–1421. 10.1016/j.celrep.2015.01.059.

